# Deconvolution of the mechanisms of T cell drug response in multiple myeloma induction therapy

**DOI:** 10.64898/2026.09.15.751861

**Authors:** Lucas T. Graybuck, Lauren Y. Okada, Wei-Ling Chang, Jessica Garber, Catalina Sakai, Morgan D.A. Weiss, Samir Rachid Zaim, Veronica Hernandez, Ziyuan He, Upaasana Krishnan, Padmapriyadarshini Ravisankar, Julian Reading, Ernest M. Coffey, Thomas F. Bumol, Jimena Meladze, Mackenzie S. Kopp, Sandra B. Munro, Gregory L. Szeto, Evan W. Newell, Xiao-jun Li, Peter J. Skene, Troy R. Torgerson

## Abstract

Lenalidomide (Revlimid™), bortezomib (Velcade™), and dexamethasone, are used alone or in combination (RVd therapy) as first-line therapies for hematologic malignancies including the plasma cell dyscrasia, Multiple Myeloma (MM). The effects of RVd treatment on tumor cells have been documented, but little is known about their impacts on “healthy” immune cells. New therapeutics, including chimeric antigen receptor T cell (CAR-T) and bispecific T-cell engagers (BiTEs) depend on robust T cell function but these are often harvested for cellular therapies after or in tandem with RVd therapy. Understanding the molecular effects of these drugs on healthy T cells is therefore of particular importance. Since the molecular effects of each drug includes regulation of key transcription factors, we used a tri-modal, single-cell assay (TEA-seq), simultaneously profiling mRNA transcripts, cell surface proteins, and chromatin accessibility in primary human peripheral blood T cells treated for 4, 24, or 72 hours *in vitro*. Synergies and conflicts potentially arising from combinatorial therapy were identified and validated by flow cytometry. Doublet and triplet drug combinations were used to assess effects on cell-surface marker expression and on T cell activation. Our results suggest that T cell function can be optimized by the administration of these drugs individually and that sequential administration may offer an opportunity to enhance T cell function by affecting localization and altering T cell states. This work provides those interested in immunology, oncology, and cell therapies an opportunity to explore molecular pathways that could be manipulated by these commonly used drugs to modify T cell function.

**One Sentence Summary:** Primary human T cells treated with multiple myeloma RVd induction therapy drugs exhibit transcriptomic, epigenetic, and phenotypic changes that should be considered and potentially leveraged in the era of T cell-dependent treatments.

## Introduction

Multiple myeloma (MM) arises from aggressive clonal expansion of plasma cells in the bone marrow (1) and is characterized by complex interactions with immune cells in the tumor microenvironment (2–4). The prognosis for MM has improved due to the addition of proteasome inhibitors and immunomodulatory drugs (IMiDs, (5)) to treatment regimens. Combination therapies comprised of proteasome inhibitor, such as bortezomib, and glucocorticoids, are the standard of care for MM induction therapy (6). Addition of the IMiD lenalidomide to this regimen has resulted in high rates of response (7). The combination of lenalidominide (Revlimid™), bortezomib (Velcade™), and dexamethasone (RVd) is highly effective as first-line treatment of newly diagnosed MM, and has demonstrated efficacy in the relapsed/refractory setting as well (8). RVd therapy is also increasingly employed as maintenance therapy for MM patients after autologous stem cell transplant (ASCT), superseding the use of paired lenalidomide and dexamethasone maintenance therapy (9). Relapse is a major concern in MM, as the acquisition of new mutations causes emergence of treatment-resistant tumor cells (10). Treatment options for relapsed/refractory MM (RRMM) have therefore expanded to include antibody (11) and T cell dependent cellular therapies including bispecific T-cell engagers (BiTEs, (12)) and chimeric antigen receptor T cells (CAR-T (13)).

Previous studies of RVd response focused largely on effects of treatment on tumor cells, but less on effects of RVd therapy on the non-malignant peripheral immune system. Of particular interest is the effect of RVd on T cells, which are essential to the generation of CAR-T or response to BiTE therapies. Multiple rounds of RVd are known to affect T cell composition, metabolism, and function in treated MM (14–16), yet there remains a paucity of information on the interplay of mechanisms underlying RVd combination treatment. Furthermore, there is little understanding of the potential synergies and conflicts that may arise in RVd combinatorial drug administration.

Each RVd drug has a unique but functionally overlapping mechanism of action making it difficult to predict their combinatorial effect. Lenalidomide binds the E3 Ubiquitin ligase Cereblon (encoded by the *CRBN* gene), resulting in proteasomal degradation of the transcription factors Ikaros (*IKZF1*) and Aiolos (*IKZF3*) that are expressed in T cells, MM tumor cells, and other cell types (17,18). In contrast, Bortezomib binds directly to the catalytic site of the 26S proteasome complex, interfering with protein homeostasis, metabolism, and regulation of multiple transcription factors including NF-kB, Ikaros, and others. Shifts in metabolic state are known to dramatically affect T cell activation, migration, and proliferation in response to stimuli (19). Dexamethasone is a glucocorticoid receptor (GR) agonist that binds directly to the GR transcription factor encoded by the *NR3C1* gene. Frequently used for its anti-inflammatory effects, Dexamethasone can abrogate the effects of immunotherapy by suppressing T cell proliferation and differentiation (20) as glucocorticoid hormone signaling affects multiple aspects of T cell biology and function in vivo (21).

We performed *in vitro* studies on the effects of lenalidomide, bortezomib, and dexamethasone on T cells isolated from healthy adults (age 31 to 62). Since each RVd drug affects at least one functionally important transcription factor family, we utilized trimodal single-cell analysis of mRNA <u>T</u>ranscripts, surface protein <u>E</u>pitopes, and chromatin <u>A</u>ccessibility (TEA-seq), a platform that permits simultaneous single-cell analysis of the cell surface proteome, transcriptome, and epigenome (22), and provides detailed resolution of the complex heterogeneity among T cells. We also tested single-, double- and triple-drug administration of RVd drugs to assess their individual and combined effects on specific cell surface markers and T cell receptor (TCR) signaling. This analysis details the effects of mono and combination therapy on T cell biology and function, providing new insight into how these drugs could be temporally administered to optimize MM treatment.

## Results

### In vitro assessment of T cell responses to drug treatment

To examine the effect of each RVd drug on T cells *in vitro*, we performed TEA-seq on primary T cells isolated from a healthy female donor aged 58 years (table S1). CD3+ T cells were isolated from cryopreserved peripheral blood mononuclear cells (PBMCs) by negative selection (Fig. 1A).

**Figure 1.**
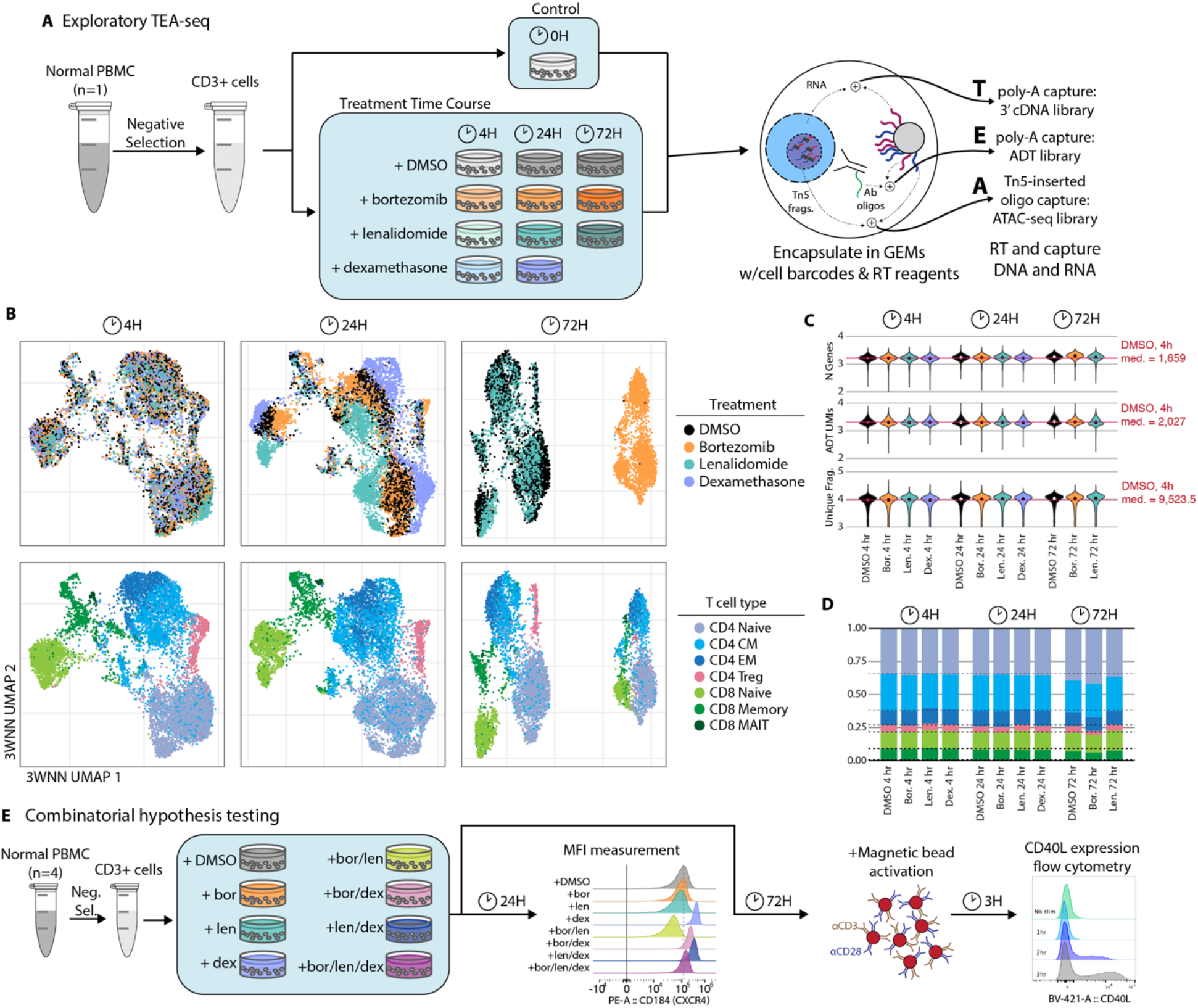
Overview of TEA-seq discovery experiment and validation experiments. **A.** Workflow diagram for drug perturbation TEA-seq experiment. **B.** UMAP projections of cells treated with DMSO only or DMSO with drug at each time point (4 hr, 24 hr, or 72 hr). Top plots are colored according to drug, bottom plots according to cell type. For display, 2500 cells per condition were randomly selected. **C.** Violin plots showing distribution of QC metrics for each modality show consistent data quality across treatment conditions. Violins are colored according to treatment, as in B. **D.** Cell-type label frequencies. Bars are colored according to cell type, as in B. **E.** Workflow diagram for combinatorial drug treatment validation experiments.

Based on pharmacokinetic reports of *in vivo* plasma concentrations for each drug, we determined appropriate concentrations of and homeostatic culture conditions to avoid causing widespread activation or cell death (fig. S1 and note S1) (23). We determined that 1000 nM lenalidomide, 2.5 nM bortezomib, and 100 nM dexamethasone were near physiological concentrations and induced drug-dependent responses over the course of 72 hours (fig. S2).

After treatment, cells were profiled using the tri-modal single cell method TEA-seq (Fig. 1A-D) as a discovery platform to deeply characterize the effect of individual drugs, followed by more specific analyses of functional effects and drug combinations (Fig. 1E).We obtained 143,828 high-quality tri-modal T cell profiles for downstream analysis (fig. S3). Data quality for all 3 modalities was consistently high across all treatments and time points (Fig. 1C). All three modalities were used for joint dimensionality reduction using the Weighted Nearest Neighbors (WNN) approach implemented by Seurat V4 (24) to provide an overview of the dataset (Fig. 1B). Cells treated with each drug exhibited a high degree of overlap at the 4-hour time point but separated by 24-hours. At 72 hours, cells treated with bortezomib distinctly separated from the other treatments.

T cell subsets were defined using epitope expression to separate CD4 and CD8 T cells, followed by independent clustering of each class and labeling of clusters based on marker epitope and gene expression (fig. S3 and Methods). We identified 7 major T cell populations for each time point: CD4 naïve, central memory (CM), effector memory (EM), and regulatory T (Treg) cells; CD8 naïve, memory, and mucosal-associated-invariant T (MAIT) cells (Fig 1B, table S2 and note S2). CD8 MAIT cells were excluded from differential analyses due to low cell abundance. Over 72 hours, frequency of cell-type populations remained consistent across treatment conditions (Fig. 1D). All differential analyses of drug response were compared to matched DMSO-treated cells to capture cell culture effects, including differentially expressed genes (DEGs, table S3), differentially detected epitopes (DDEs, table S4), and differentially accessible ATAC peaks (DAPs, table S5).

### Lenalidomide degrades Ikaros and Aiolos

Previous studies have shown that the IMiD Lenalidomide causes T cells to develop effector phenotypes and functions, potentially enhancing their activity against tumor cells and rejuvenating exhausted CD8+ T cells to an effector state (25,26). It is hypothesized these effects may be through Lenalidomide’s regulation of the E3 ubiquitin ligase Cereblon that mediates ubiquitination and proteasomal degradation of Ikaros family transcription factors (17).

Intracellular flow cytometry on resting and activated T cells treated with lenalidomide confirmed that it induced degradation of Ikaros (encoded by the *IKZF1* gene) and Aiolos (*IKZF3*) proteins in our *ex vivo* T cell culture system (Fig. 2A-B). While Ikaros and Aiolos levels remained low in resting, lenalidomide-treated T cells, activation increased Ikaros levels back to the untreated baseline level (Fig. 2A-B). However, Aiolos was initially up-regulated, then degraded in both drug-treated and untreated cells following activation. In our TEA-seq data, we found that *IKZF1* gene expression was significantly up-regulated in most T cell types, likely to compensate for the loss of Ikaros protein (Fig. 2C). *IKZF1* was detected in fewer cells and at lower levels in Treg and CD8 memory cells without treatment, though CD8 memory cells responded to lenalidomide by up-regulating *IKZF1* transcription. We examined the accessible chromatin landscape near the *IKZF1* gene in the same cells and identified a region of differentially accessible chromatin ∼75 kb upstream of the *IKZF1* transcription start site (Fig. 2D). Lenalidomide affected the accessibility of *IKZF1* chromatin in cell types with compensatory *IKZF1* gene expression, but not in Treg cells, which showed no compensatory *IKZF1* up-regulation (Fig. 2D). Analysis of transcription factor (TF) binding sites in this region revealed multiple sites for TFs expressed in T cells, including ATF4, NFATC1, NFATC2, NFATC3, STAT1, STAT3, and STAT5B (27,28).

**Figure 2.**
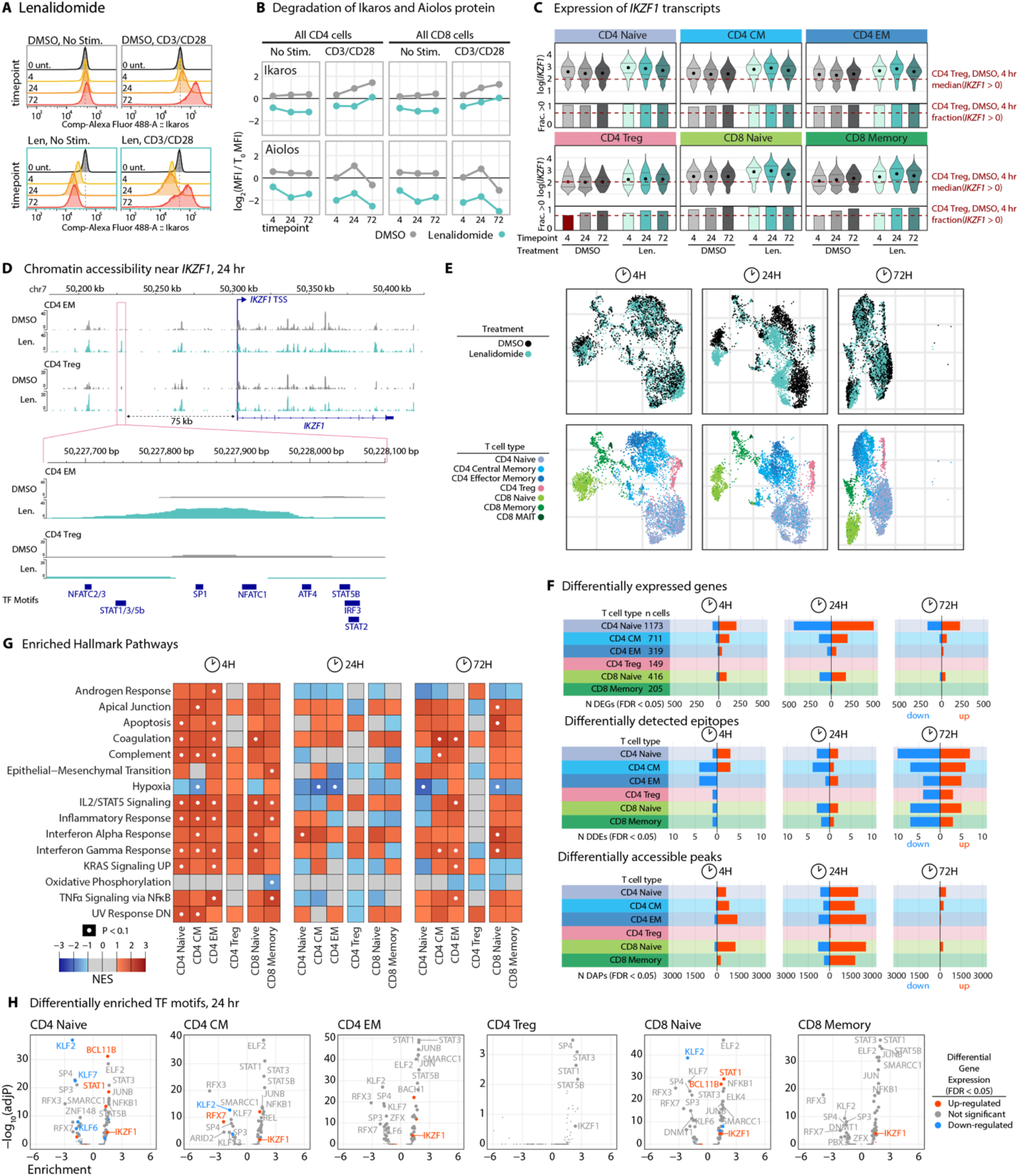
Effects of lenalidomide on T cell state. **A.** Representative histograms for changes to Ikaros by intracellular flow cytometry of CD4 T cells. Untreated cells (black) are compared to cells treated for 4 (yellow), 24 (orange), or 72 hours (red). Dashed line represents the mode of untreated control. Y-axes are normalized to the mode of each condition. X-axes use a log-transformed scale of compensated values. Top row, DMSO only; bottom row, Lenalidomide; left panels, Unstimulated; right panels, Stimulated; Unt., untreated. **B.** Mean fluorescence intensity (MFI) changes to Ikaros/IKZF1 and Aiolos/IKZF3 protein expression. Line plots show MFI changes relative to untreated T cells on a log_2_ scale. DMSO only, gray lines; Lenalidomide, green lines. **C.** Violin plots of normalized expression *IKZF1* in each cell type. Violin plot values are drawn from non-zero expression values. Points represent medians; lines represent the 25th and 75th percentiles. Barplots show the fraction of cells with non-zero expression (fraction detected). Dashed red lines show median expression and fraction detected for CD4 Treg cells. **D.** Chromatin accessibility near the IKZF1 gene for CD4 EM and CD4 Treg cells at 24 hours. Y-axis, scATAC-seq fragment overlap. Solid bars, motif locationsCoordinates are from the GRCh38 genome assembly. **E.** UMAP projections highlighting DMSO and lenalidomide treated cells at each timepoint. **F.** Barplots of up-regulated (red) and down-regulated (blue) DEGs (top panel), DDEs (middle panel), and DAPs (bottom) at each time point. **G.** Top Hallmark pathway Gene Set Enrichment scores as shown in Figure 2B for lenalidomide. **H.** Volcano plots for transcription factor motif enrichment as in Figure 2D for lenalidomide treatment at 24 hours.

Next, we examined the TEA-seq profiles of lenalidomide response over time and identified substantial but transient changes to T cell gene expression (tables S3, S4, and S5). Lenalidomide induced rapid changes in gene expression by 4 hours, increasing by 24, and subsiding by 72 (Fig. 2E-F). We used gene set enrichment analysis (GSEA) to test for enrichment of gene sets among differential expression results. We compared DEG for each time point to the Molecular Signatures Database (MsigDB) Hallmark gene sets (29) (Fig. 2C and table S6) and the Reactome Pathways database (30) (fig S4, table S7). Lenalidomide treatment resulted in up-regulation of genes in Hallmark pathways related to apoptosis, inflammation, and interferon response, MYC targets, and TNF-α signaling (Fig. 2G and fig. S4A). Despite more observed DEGs at 24 hours, Hallmark pathway enrichment results were more strongly observed at 4 hours post-treatment than at 24 hours. A similar pattern was observed in chromatin accessibility, with an initial wave of increased accessibility at 4 hours, a substantial increase at 24 hours, followed by less differential accessibility at 72 hours (Fig. 2F). As observed in *IKZF1* transcription, Treg cells showed fewer changes in expression or accessibility than other types (Fig. 2F).

In our multimodal dataset, we found that lenalidomide induced differential accessibility associated with multiple transcription factors, including increased accessibility at IKZF1, STAT5B, and NFKB1 motifs. IKZF1 has been previously shown to control access to NFKB1 motifs (31). However, this increase in IKZF1-related site accessibility does not occur in Treg cells (Fig. 4H). As with transcriptional changes, changes to chromatin accessibility were also largely transient, and decreased by 72 hours relative to their peak at 24 hours after lenalidomide administration (Fig. 4F).

### Bortezomib disrupts protein homeostasis and suppresses T cell receptor-related gene expression

Bortezomib is a boron modified di-peptide that inhibits the proteosome complex. In T cells it alters gene expression and chromatin accessibility at 24 and 72 hours to cause an increasingly perturbed T cell state (Fig. 3A and B, tables S3, S4, and S5). The changes in chromatin accessibility led to enrichment of transcription factor motifs important for bortezomib response (Fig. 3D).

**Figure 3.**
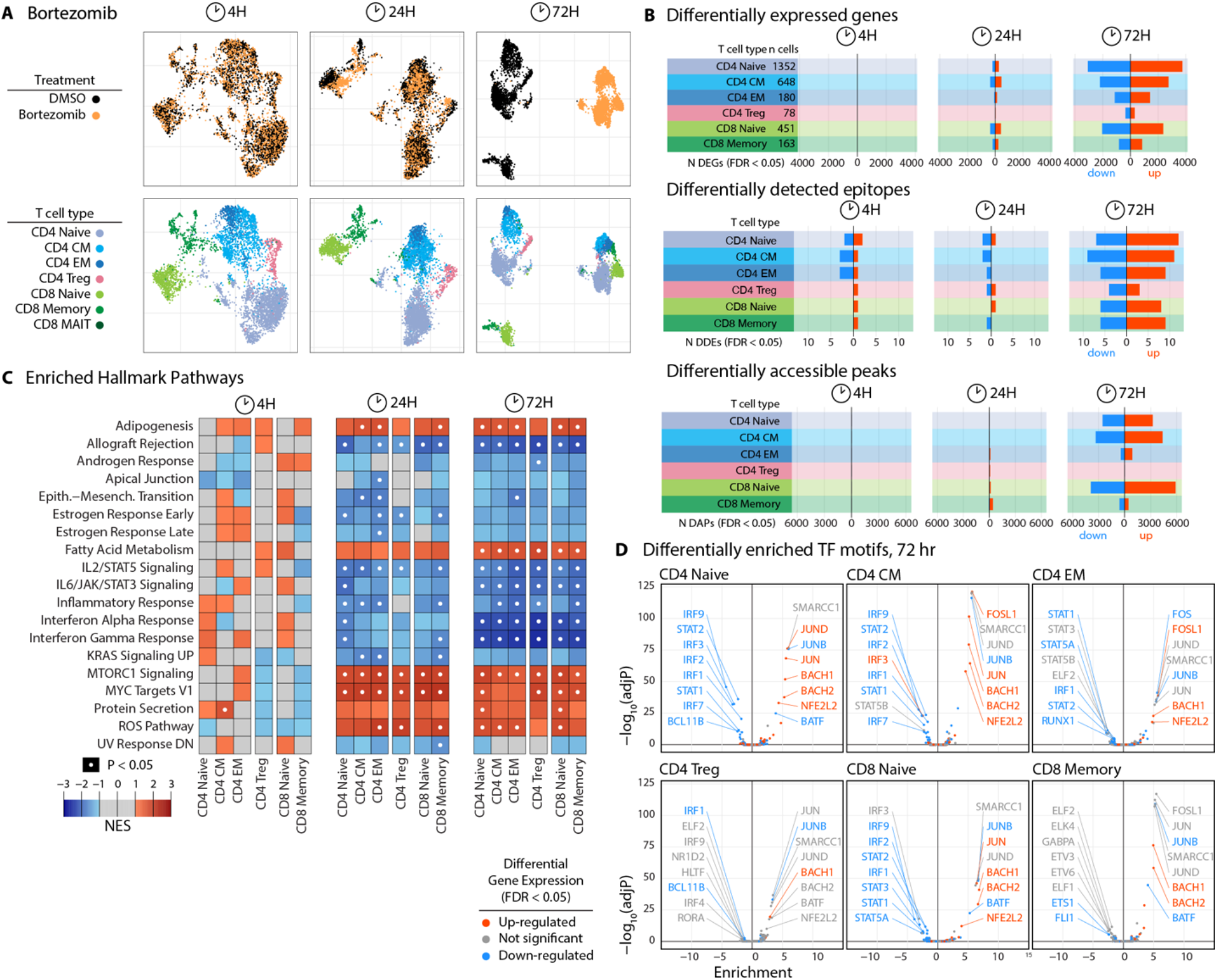
Effects of bortezomib on T cell state. **A.** UMAP projections highlighting DMSO and bortezomib treated cells at each timepoint. **B.** Barplots showing the number of up-regulated (red) and down-regulated (blue) genes (top panel), cell surface epitopes (middle panel), and accessible chromatin regions (bottom) at each time point. **C.** Top Hallmark pathway Gene Set Enrichment scores for differentially expressed genes. Each row represents a Hallmark pathway, and each column represents a cell type. Red, up-regulated genes; blue, down-regulated genes. Pathways with significant enrichment indicated with a white dot (FDR < 0.1). **D.** Enriched transcription factor motifs identified in differentially accessible peaks at 72 hours after treatment. Each panel presents enrichments for a cell type at 72 hours. Points are colored based on differential gene expression of the transcription factor that binds to each motif: red, up-regulated; blue, down-regulated; gray, no significant change (FDR values < 0.05 considered significant.).

The MTORC1 Signaling and MYC Targets V1 Hallmark gene sets were among the most strongly and consistently up-regulated pathways in T cells treated with bortezomib for 24 and 72 hours (Fig. 3C). These pathways include several genes encoding proteosome subunits, including *PSMA3*, *PSMA4*, *PSMB5*, *PSMC2*, *PSMC4*, *PSMC6*, *PSMD12*, *PSMD13*, and *PSMD14*. Analysis of the transcriptional response of proteasomal and proteasome-related genes (as described in (32)) revealed widespread up-regulation across multiple proteasome subunits (Fig. 4A), apart from the immunoproteasome and PA28/11S components. Previously identified targets of bortezomib are well-represented in this gene set, including *PSMB5,* the gene for the active site protein in the 26S proteasome, a common site of mutations that confer bortezomib resistance (33).

**Figure 4.**
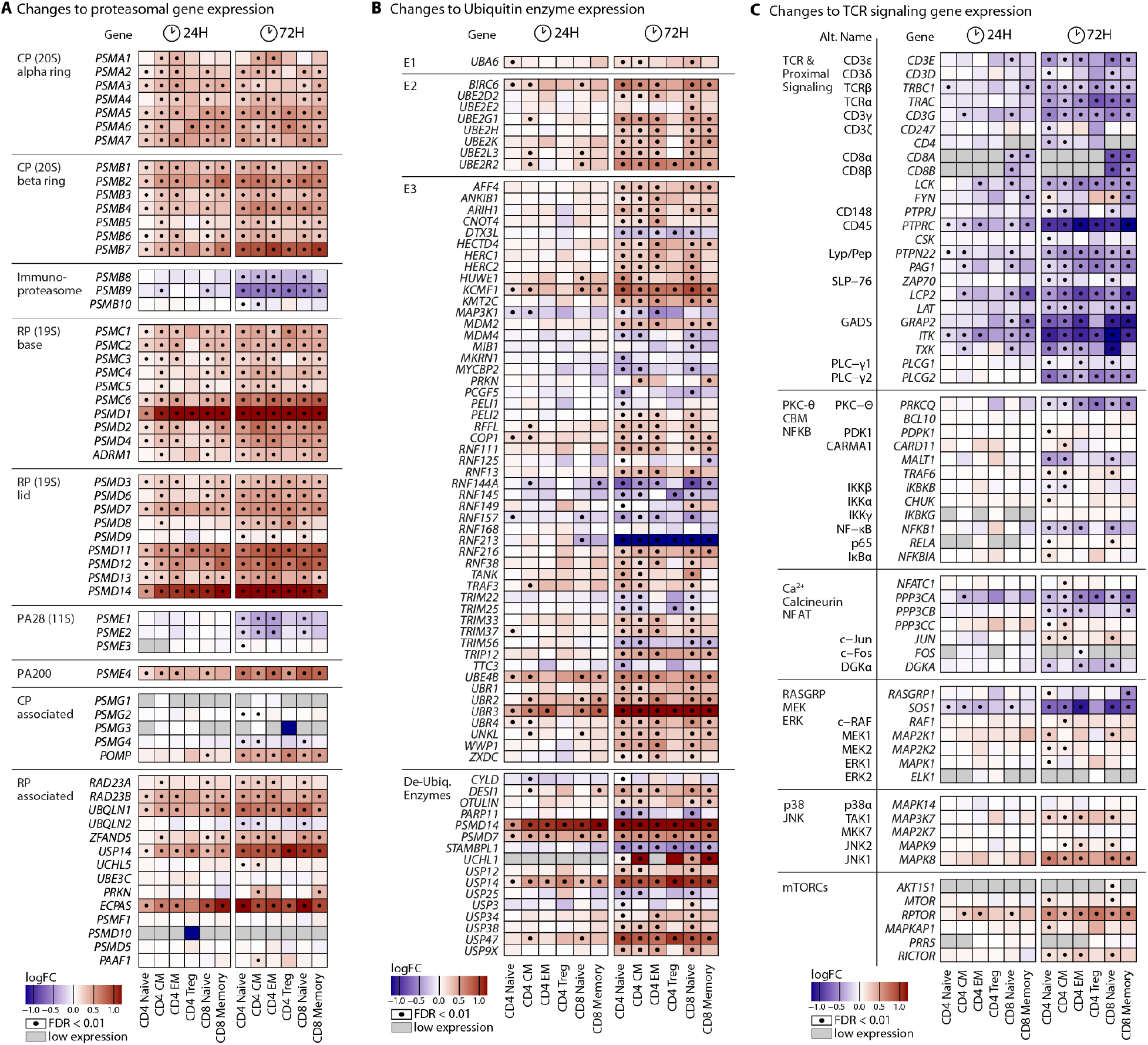
Differential expression of proteasomal, ubiquitin, and TCR signaling genes. **A.** Heatmap displaying the Log Fold Change in gene expression for proteasomal genes described in (32). Each row represents one gene, and each column represents differential expression for a cell type (bottom) at the 24 hr or 72 hr time point (top). Significant DEGs are indicated with a black point (FDR < 0.01). Gray box, expression too low to test. CP, Core Particle; RP, Regulatory Particle; PA, Proteasome Activator. **B.** Heatmap as in A. for ubiquitination-related protein genes collated in the HGNC Gene Groups (E1 and E2, (66)) or the ESBL (all others, (67)). Only differential genes in at least one comparison are displayed (FDR < 0.01 and logFC > 0.3). E1, Ubiquitin activating enzymes; E2, Ubiquitin conjugating enzymes; E3, Ubiquitin ligase enzymes; De-Ubiq, De-ubiquitinating. **C.** Heatmap as in A. for genes in TCR activation and downstream processes, as described in (68).

The proteasome associated deubiquitinase genes *PSMD7* and *PSMD14* were highly upregulated across multiple subsets at the 24 and 72 hour timepoints. To define whether ubiquitylation processes were broadly disrupted by bortezomib, we examined changes in expression of a panel of ubiquitin-related genes (Fig. 4B). *DESI1*, *USP14* and *USP47* deubiquitinase genes were up-regulated at 72 hours along with several other ubiquitin-related genes that were up-regulated in response to bortezomib (Fig. 4B). *OTULIN*, a deubiquitinase for linear ubiquitin chains that plays a role in regulating inflammatory responses is among these (34).

When proteasome-related genes are removed from pathway enrichment tests, the MYC Targets V1 gene set is no longer significantly enriched, indicating that the overlap with this pathway is primarily due to proteasomal gene up-regulation (fig. S6). However, down-regulation of inflammatory response pathways, including those of IL2/STAT5 signaling, IL6/JAK/STAT3 Signaling, Inflammatory Response, Interferon Alpha, and Interferon Gamma response remained significant (fig. S6). These pathways include several inflammation-related genes including *IL4R*, *IRF1*, *PIM1*, *TNFRSF1B*, *GBP4*, *LY6E*, *SOCS1*, *SELL*, and *TNFSF10* with reduced expression across multiple T cell subsets.

Several inflammatory response pathways important for T cell signaling such as NF-κB activation, require ubiquitination and proteasome degradation of inhibitory proteins (35,36). We therefore examined differential expression of genes involved in TCR activation and downstream signaling pathways (Fig. 4C) that are key to T cell activation but are not well described by Hallmark gene sets. We found that in all T cell subsets, bortezomib caused down-regulation of many genes in the TCR complex and in TCR-proximal signaling (Fig. 4C). In downstream signaling pathways, we found down-regulation of PKC-Θ (*PRKCQ*) and *MALT1*, a member of the CARD-BCL10-MALT1 (CBM) complex. We observed down-regulation of NF-κB (*NFKB1*), a key transcription factor for activation of gene expression in response to T cell activation, and Son of Sevenless (*SOS1*), a Ras-specific guanine-nucleotide exchange factor (37). We also observed up-regulation of genes in the JNK kinase cascade: TAK1 (*MAP3K7*) and JNK1 (*MAPK8*), as well as genes in mTOR complexes: *RPTOR* and *RICTOR*.

In addition to transcriptional changes, TEA-seq revealed widespread changes in chromatin accessibility in response to bortezomib at 72 hours (Fig. 3B). Among sites with increased accessibility, ATF4, ATF7, BACH1, BACH2, JUN, JUNB, MAFG, NFE2L2, RUNX1, and SMARCC1 motifs were enriched across all T cell subsets (Fig. 3D, table S8). BACH- and MAF-family proteins are known to directly interact to regulate gene expression, and can be involved in both epigenetic silencing and up-regulation of target genes depending on other protein interactions (38). In sites with decreased accessibility, ELK-family, IRF-family and STAT-family transcription factor motifs were enriched among all T cell types except Tregs (Fig. 3D, table S8).

We also observed changes to cell surface protein expression in response to bortezomib (Fig. 3B and fig. S7). The most pronounced changes were observed at 72 hr, including up-regulation of CD69, CD127 (encoded by the *IL7R* gene), CD38, CD95 (*FAS*), and CD278 (*ICOS*) expression and down-regulation of CD71 (*TFRC*), CD27, CD196 (*CCR6*), and CD269 (*PDCD1*). Some of these changes countered effects observed over time in treatment with DMSO alone (fig. S7), which may be because of the IL-2, IL-7, and IL-15 cytokine cocktail used for all experiments. For example, the IL7 receptor (CD127), is strongly down-regulated in DMSO treated control cells compared to freshly-thawed cells. This trend is reversed with bortezomib treatment at 72 hours (fig. S7).

### Dexamethasone induces tissue homing- and retention-related genes

The glucocorticoid drug dexamethasone is widely used for its broad anti-inflammatory function. In multiple myeloma RVd therapy, it reduces inflammation, swelling, and pain at sites of tumor invasion in tissues but also directly causes apoptosis of myeloma cells and sensitizes them to the effects of the other drugs (39). To understand how dexamethasone affects T cells, we utilized TEA-seq and observed a rapid response at 4-hours of treatment with an expanded response at 24 hours (Fig. 5A-B).

**Figure 5.**
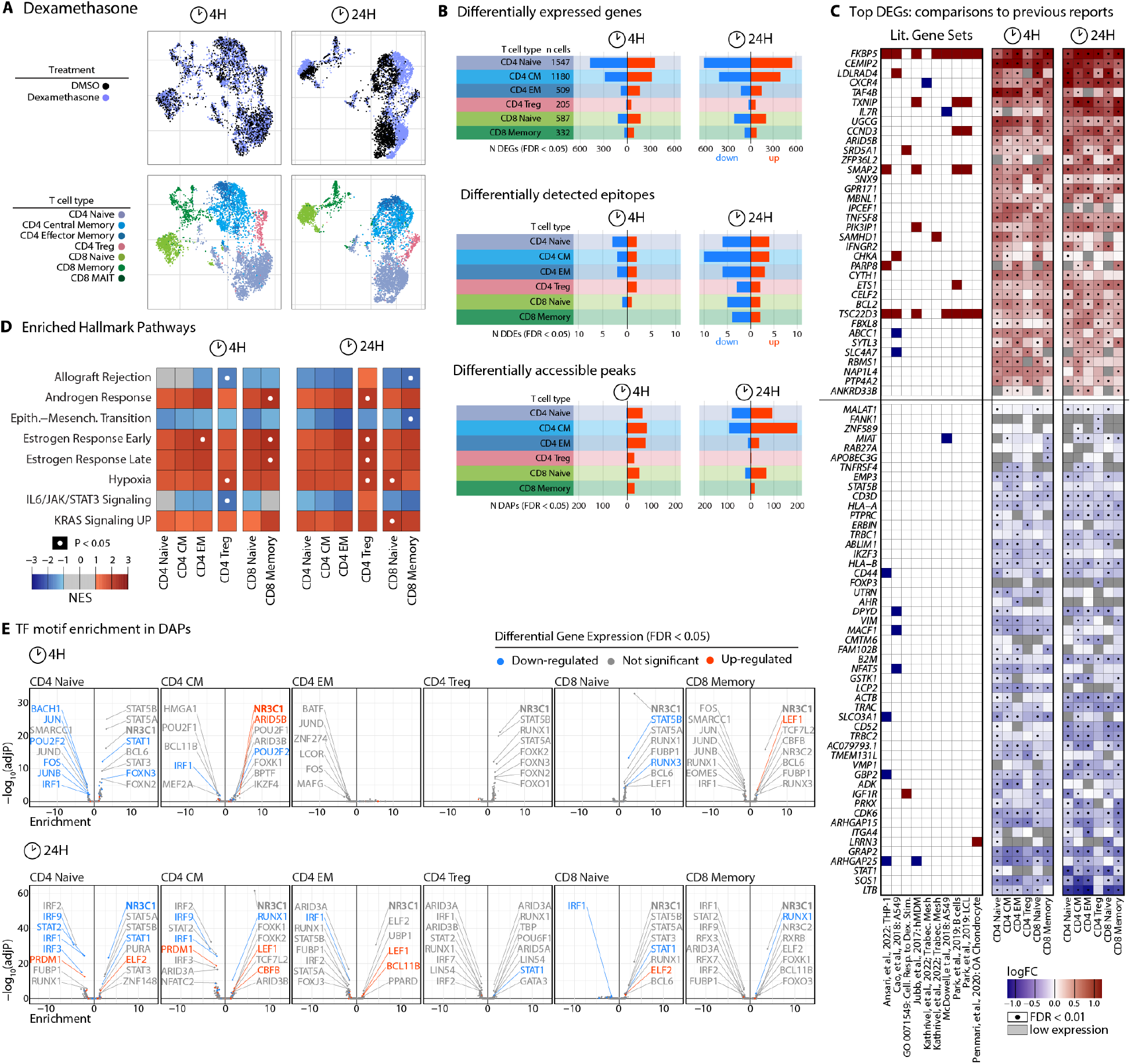
Effects of dexamethasone on T cell state. **A.** UMAP projections highlighting DMSO and dexamethasone at 4 hr and 24 hr time points. **B.** Barplots of up-regulated (red) and down-regulated (blue) DEGs (top panel), DDEs (middle panel), and DAPs (bottom) at each time point. **C.** Heatmap of the most highly differential genes at 4 hr and 24 hr after dexamethasone treatment and occurrence in previous reports. Each row represents a gene. In the left panel, each column represents a published dataset (table S8). Genes that are also among the most differentially expressed in our T cell gene set are colored: up-regulated, red; down-regulated, blue. **D.** Top Hallmark pathway Gene Set Enrichment scores as in Figure 2B for dexamethasone. **E.** Enriched transcription factor motifs identified in DAPs for T cell subsets after 4 hours (top) and 24 hours (bottom).

We surveyed the literature for datasets that captured responses to dexamethasone in primary human cells or cell lines (table S9) and compared these gene sets to the top DEGs observed in our TEA-seq data (table S3). The top 20 up-regulated and top 20 down-regulated DEGs for each cell type were selected, resulting in a set of 37 unique up-regulated and 44 unique down-regulated genes. Of these, only 17 (39%) up-regulated and 5 (11%) down-regulated genes were identified in these previously published datasets (Fig. 5C). Many genes not represented in previously published datasets are important for T cell function: cell surface receptors are up-regulated, including *CXCR4*, the receptor for the chemokine CXCL12, *IL7R*, the receptor for the cytokine IL-7, *IFNGR2*, encoding the beta chain of the interferon gamma receptor, and *TNFSF8*, encoding the ligand for the CD30 receptor. Conversely, *TNFRSF4*, encoding the receptor for the co-stimulatory OX40-ligand molecule, is down-regulated in response to dexamethasone, as are major histocompatibility complex I genes *HLA-A*, *HLA-B*, and *B2M. PTPRC*, which encodes the CD45 component of the TCR signaling complex, is also downregulated.

Hallmark pathways related to androgen and estrogen responses, whose receptors share targets with the glucocorticoid receptor (40), were enriched among up-regulated genes in CD4 EM, Treg, and CD8 Memory cells at 24 hours post-treatment (Fig. 5D). At 24 hours, pathways related to hypoxia were upregulated in CD4 EM, Treg, CD8 naïve, and CD8 memory cells (Fig. 5D). In REACTOME gene sets, we also observed immunosuppressive signatures: association of down-regulated genes with TCR signaling, CD28 costimulatory signaling, and Class I MHC signaling across most T cell types, as well as Interferon signaling pathways in CD4 EM and CD8 Memory cell types (fig. S4B).

We observed an initial increase of chromatin accessibility at 4 hours, which expanded at 24 hours to include regions of decreased accessibility relative to DMSO controls (Fig. 5B). We observed fewer changes in accessibility in Tregs and CD8 Memory cells compared to other T cell types. In CD4 EM cells, we saw enrichment of NR3C1/NR3C2, STAT5B, STAT1, RUNX1, and FOX-family transcript motifs among peaks with increased accessibility at 4 hours post-treatment, and JUN/FOS, BACH1/2, and IRF2 motifs enriched among the few peaks with decreasing accessibility at 4 hours (Fig. 5E). At 24 hours we saw additional secondary responses that included RUNX1, STAT2, PRDM1, and IRF1/2 motif enrichment among sites with decreased accessibility, and ZNF148, STAT5A/B, and FOXK1/2 motifs enriched in peaks with increased accessibility in multiple cell types (Fig. 5E).

### Identification of potential drug interactions

Multiple Hallmark pathways were enriched in opposite directions by different drugs: inflammatory and interferon responses were up-regulated by lenalidomide, but down-regulated by bortezomib, and MYC targets were up-regulated by bortezomib, but down-regulated by dexamethasone. To identify potential drug synergies and conflicts at the gene level, we compared DEGs from each drug within each T cell subset(Fig. 6A and fig. S8). Overlaps between differential sets were tested for significant enrichment using hypergeometric tests (fig. S8 and table S10).

**Figure 6.**
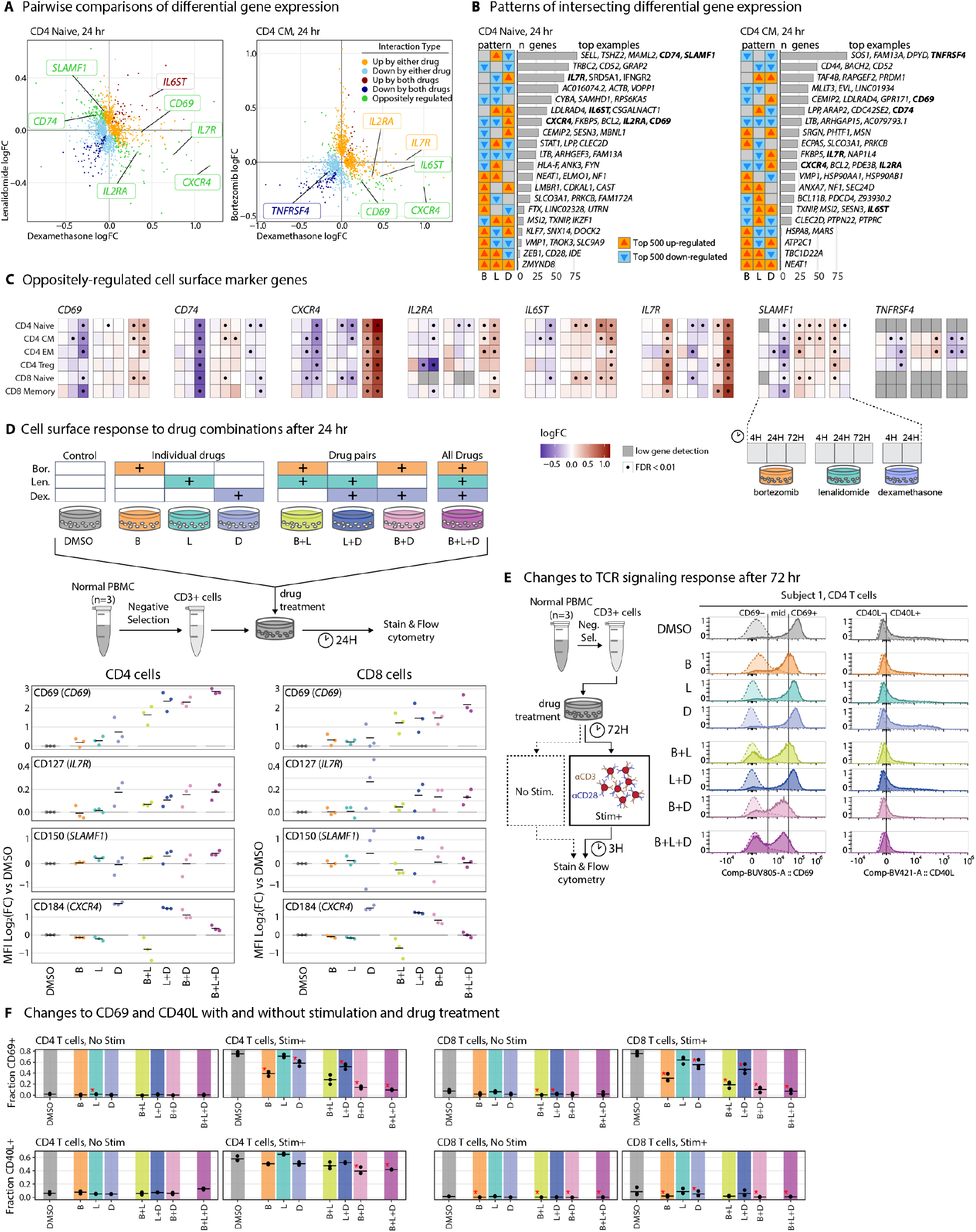
Effects of drug co-administration on T cell surface epitopes and function. **A.** Pairwise comparisons of between two different drug treatment conditions at 24 hours. Points are individual genes, and are colored based on agreement or disagreement between the drug conditions: orange, up-regulated by one drug; light blue, down-regulated by one drug; dark red, up by both drugs; dark blue, down by both drugs; green, oppositely regulated. **B.** Barplots showing the number of differential genes with each pattern of differential expression. Arrows indicate whether the set was up- or down-regulated by each drug. **C.** Heatmaps for change in transcription of selected cell surface markers. Each row represents a cell type. Each gene is represented by a set of columns that show the comparisons between drug and DMSO at each time point, as shown in the callout below the *SLAMF1* panel. Genes with FDR < 0.01 are indicated with a black point. **D.** Selected cell surface marker expression measured by flow cytometry following treatment with individual drugs, pairs of drugs, or all 3 drugs for 24 hours, as diagramed in the top panel. Each point represents log fold change in MFI from a single donor (N = 3). Drug conditions are separated along the x axis, and points are colored based on the drug combination legend in the top panel. **E.** Histograms of key TCR signaling response proteins as measured by flow cytometry following treatment of T cells with each drug combination, as outlined in (**C**). **F.** Fractions of CD69+ (top) or CD40L+ (bottom) cells in CD4 (left) and CD8 (right) T cells with and without stimulation, as shown in (**E**). Each point is a replicate. Bars show medians, and a red asterix indiciates an adjusted P-value < 0.05 in comparisons to the DMSO control.

We systematically identified genes whose expression changed in opposite directions by any pair of the 3 drugs (Fig. 6B and table S10), then filtered the results for known cell surface epitopes to identify candidates for validation by flow cytometry. We selected a panel of 8 targets that represented a variety of conflict patterns (Fig. 6C): CD69, CD74, CXCR4 (CD184), IL2RA (CD25), IL6ST (CD130), IL7R (CD127), SLAMF1 (CD150), and TNFRSF4 (CD134). We tested co-administration of drugs by treating healthy T cells from 3 different donors with no drug, each individual drug, each drug pair, and all 3 drugs together for 24 hours, followed by surface staining for these 8 marker proteins or related complexes (HLA-DR in the case of CD74), and additional markers that enabled identification of CD4 and CD8 T cell populations (fig. S9 and table S11).

For 4 of the 8 targets, we saw changes in cell surface expression across individuals (Fig. 6D and fig. S9). We observed increased CD69 expression by flow cytometry in response to dexamethasone treatment, as observed transcriptionally. Despite down-regulation of *CD69* gene transcription by bortezomib and little effect by lenalidomide, we saw an additive effect of administration of each pair of drugs or all 3 drugs together (Fig. 6D). IL7R was transcriptionally up-regulated by both bortezomib and dexamethasone but down-regulated by lenalidomide in some T cell subsets. In flow, we observed an increase with dexamethasone and any combination treatment that included dexamethasone. *CXCR4* gene expression and CD184 protein expression were strongly increased by dexamethasone treatment, as previously reported (41,42). Treatment with both bortezomib and lenalidomide caused down-regulation of *CXCR4* expression and a modest decrease in CD184, and combinations of either drug with dexamethasone reduced CD184 protein expression from the highly induced level driven by dexamethasone alone. Co-administration of all 3 drugs either antagonized the dexamethasone-induced increase in CD184 protein (in CD4 cells) or reduced it to match the effect of DMSO alone (in CD8 cells).

We observed many differences in gene expression that may affect T cell response to TCR stimulation and downstream signaling in response to individual drugs (Fig. 3C, fig. S4, fig. S5). Bortezomib induced strong down-regulation of many TCR and TCR-associated genes and downstream signaling pathways (Fig. 3C). Lenalidomide and dexamethasone both had less widespread effects on TCR signaling-related gene expression, but each induced alterations to expression of key regulatory genes in downstream pathways. To better understand how individual and combinatorial drug treatment affects functional TCR responses, we treated T cells using the combinations outlined in Fig. 6C for 72 hours, followed by stimulation with CD3/CD28 beads or no stimulation for 3 hours, then observed TCR response using CD69 and CD40L expression as markers of activation (Fig. 6F, fig. S10 and table S12). In CD4 and CD8 T cells, we found that treatment with any individual or combination of drugs decreased CD69 expression in response to TCR stimulation, with the greatest effects observed using bortezomib or combinations that included bortezomib (Fig. 6F). Coadministration of all 3 drugs had the greatest suppressive effect. Among individual drugs, lenalidomide treatment had the least suppressive effect, and among pairs of drugs, lenalidomide with dexamethasone was least suppressive. A similar pattern was observed for CD40L expression on CD4 T cells in response to TCR stimulation, but to a lesser extent – only bortezomib with dexamethasone and coadministration of all 3 drugs significantly suppressed CD40L expression compared to the DMSO control (adjusted *P*-value < 0.01, Fig. 6G).

## Discussion

Combinatorial induction therapies such as RVd therapy are effective first-line treatments for MM and other cancers that engage distinct biological processes to target vulnerabilities in tumor biology. However, long-term consequences of therapy may arise, so subsequent treatments that engage immune cells to target RRMM tumors may be affected. In a previous study, ASCT and lenalidomide/dexamethasone maintenance therapy were found to alter the balance of CD4 and CD8 T cell frequency, alter T cell metabolism, and increase expression of aging and exhaustion markers (14). Long-term RVd therapy can alter the functional landscape of the immune system, as shown in a recent study of NDMM treatment and post-ASCT recovery that found variable vaccine responses that correlated with residual defects in immune reconstitution after ASCT (43).

Since the molecular mechanism of action for each drug is known to include effects on transcription factor families important for T cell function, we used TEA-seq to perform deep multimodal characterization of each individual drug’s effect on the homeostasis and function of primary human T cells. We found that RVd drugs had effects related to T cell metabolism, effector state, tissue-homing and retention, and signaling, each of which play important roles in downstream T cell-dependent therapies. We identified potential conflicts between drug treatments that may alter responses in unexpected ways. These same responses have been identified in CAR-T studies, where the composition of CD4:CD8 (44) and memory:effector states (45) have been shown to influence efficacy, and studies of BiTE therapy, where T cell health prior to therapy is important for clinical response (15).

Lenalidomide changes the targeting of the E3 ligase Cereblon to cause selective ubiquitylation and degradation of the Ikaros and Aiolos zinc finger transcription factors (17). Previous studies have demonstrated increases in chromatin accessibility at sites containing IKZF1 motifs following degradation of Ikaros, supporting a repressive role for IKZF1 in chromatin regulation (46,47). We observed degradation of Ikaros protein in T cells treated with Lenalidomide (Fig. 2A-B), increased accessibility of IKZF1 motifs in chromatin (Fig. 2H), and compensatory up-regulation of *IKZF1* gene transcription over 72 hours (Fig. 2C). In addition, we identified a differentially accessible region that may contribute to compensatory up-regulation of *IKZF1* gene transcription that was only utilized in non-Treg cell types (Fig. 2D). As shown in previous mouse studies (48), loss of Ikaros results in increased accessibility of NFKB1 and STAT5B motifs. In our TEA-seq data, these changes were short-lived and resolved by the 72-hour time point (Fig. 2F). However, lenalidomide is frequently administered as a daily oral medication during induction and maintenance therapies for MM. Recurrent exposure may maintain T cells in a lenalidomide-induced epigenetic state *in vivo* may contribute to the changes in T cell homeostasis and aging observed in patients on lenalidomide maintenance therapy (14). Lenalidomide-induced changes could also have positive effects in the setting of CAR-T activity, where lenalidomide has been shown to advance an NF-κB-driven pro-effector state to enhance anti-BMCA CAR-T activity and delay exhaustion (49).

Among the 3 RVd drugs studied, Bortezomib had the largest effect on transcription and chromatin accessibility, (Fig 3) triggering widespread compensatory transcription of proteasome subunit genes (Fig. 4A). We also found widespread changes to ubiquitination machinery, including increases in proteasomal and non-proteosomal de-ubiquitination as well as up- and down-regulation of other key ubiquitin-related factors (Fig. 4B).

Bortezomib also altered cellular metabolism, leading to down-regulation of respiratory electron transport-related genes (fig. S5), which shifts T cells to use glycolysis as their primary pathway for energy generation. While this shift was observed across all T cell types, this metabolic state may favor effector T cells over naïve and central memory cell types, which rely on oxidative phosphorylation to maintain their metabolic state over long periods of time (19). In a recent study of induction therapy, ASCT, and post-ASCT recovery, a pattern of effector T cell expansion and naïve and helper T cell deficits in post-ASCT reconstitution was observed, which is consistent with this metabolic shift in T cells (43), and is suggestive of a durable metabolic shift induced by bortezomib treatment that may be better tolerated by effector cells, resulting in compositional shifts in the T cell compartment.

We also observed an increase in activation marker *CD69* in response to bortezomib (Fig. 6C). However, T cells also down-regulated many genes related to TCR signaling responses, resulting in reduced surface expression of CD40L in response to activation (Fig. 6E), and suggesting that CD69 protein expression was decoupled from T cell activation. An increase in CD69 expression on the surface of peripheral T cells was also observed in a cohort of NDMM patients undergoing RVd induction therapy, without expression of *ITGAE* (encoding CD103) or *CCR7* (CD197) that would reflect a tissue residency program (43). Previous studies have identified a role for CD69 in cellular metabolism in cooperation with the CD98/LAT1 amino acid transporter (50). In our study, the *SLC3A2* gene, which encodes CD98, was up-regulated in response to bortezomib (table S3). These data suggest that, in the context of proteasome inhibition, CD69 may be preferentially expressed as part of a metabolic response independent of T cell activation.

Unlike lenalidomide and bortezomib, that alter transcription factor functions by modulating different aspects of the ubiquitin-proteasome system (UPS), dexamethasone acts directly by binding to the glucocorticoid receptor (GR) transcription factor, encoded by the *NR3C1* gene. This results in its translocation from cytoplasm to nucleus, where it can regulate genes near GR elements (GREs)/NR3C1 motifs (41). In our data, early dexamethasone responses were enriched for previously observed estrogen and androgen receptor-driven transcriptional responses (Fig. 5D) and for many cell-type-specific changes not observed in previous human transcriptomic studies involving non-T cell types (Fig. 5C). These changes were initially driven by a robust GR response observed at the chromatin level, leading to an initial wave of increased chromatin accessibility (Fig. 5E). This was followed by reduced accessibility of regions containing IRF-family motifs, as well as reduction in accessibility of motifs for the PRDM1/BLIMP1 transcriptional repressor in CD4 naïve and CM cells at 24 hours (Fig. 5E). Glucocorticoids have been shown to up-regulate the *PRDM1* gene (51), consistent with the up-regulation of *PRDM1* at both 4 and 24 hours that we observed after dexamethasone treatment (supp. Table 3).

Among the most highly up-regulated genes were the chemokine receptor *CXCR4*, and the IL-7 receptor *IL7R* (Fig. 6C), both of which have previously been observed in diurnal T cell responses to glucocorticoids (42). CXCR4 regulates tissue homing responses and is strongly up-regulated in some MM tumors (52). A well characterized counter to CXCR4 signaling is the sphingosine-1-phosphate receptor, encoded by the *S1PR1* gene, which regulates lymphocyte egress from lymphoid organs (53). We found no up-regulation of *S1PR1* in response to dexamethasone (table S3), leaving open the possibility that dexamethasone may increase T cell migration into the bone marrow by increasing *CXCR4* expression, where they could better target MM tumor cells. This approach has been pursued in recent mouse model studies, in which electroporation of NK cells with *CXCR4* templates yielded better tissue homing of CAR-NK cells (54), treatment of CAR-T cells with dexamethasone prior to administration up-regulated IL7R and enhanced tumor killing when IL-7 was co-administered as part of therapy (55), and CXCR4 overexpression in CAR-T cells has been shown to increase efficacy of CD19 CAR-T cells in B cell lymphoma and BCMA CAR-T cells in treatment of MM (56).

Simultaneous treatment of cells with lenalidomide and bortezomib has been shown to inhibit the effects of lenalidomide by blocking degradation of Ikaros. However, if treatment with lenalidomide preceded treatment with bortezomib, degradation of Ikaros occurred prior to proteasome inhibition (57). The possibility of this and other interactions that could affect drug response and T cell function led us to compare drug treatment responses. While drug treatments were self-consistent across time – e.g., genes up-regulated at one time point significantly overlapped with genes that were up-regulated at a later timepoint (fig. S8), there were many instances of opposing changes induced by different drugs. Drug conflicts, particularly inhibition of lenalidomide and dexamethasone responses by bortezomib, emerge as a consequence of simultaneous coadministration of all 3 drugs in RVd therapy (Fig. 6 and fig. S8). For example, lenalidomide, dexamethasone, or both did not inhibit CD69 up-regulation in response to TCR activation, while treatment with bortezomib or bortezomib with dexamethasone resulted in up-regulation of CD69 regardless of activation (Fig. 6E-F), possibly due to a role of CD69 in amino acid transport as a consequence of proteasome inhibition.

Due to the broad effects of proteasome inhibition, including counteracting lenalidomide-driven Ikaros and Aiolos degradation, reducing T cell activation responses, and decreasing dexamethasone-driven CXCR4 up-regulation, these data raise the question of whether administration of dexamethasone and/or lenalidomide prior to administration of bortezomib could enhance tissue homing and retention of T cells in MM treatment. Thereby providing an avenue for improved RVd therapy. However, this hypothesis requires additional testing in model systems that better capture the state of the MM patient immune system, including hyperproteinemia (58,59) and signaling generated by, or in response to, MM tumor cells. These conditions are not well represented in our *in vitro* drug profiling experiments. In addition, treatment courses for MM are performed across many rounds of therapy, often lasting for months in total, and maintenance therapy may continue for multiple years. Here, we have profiled the effects of acute drug responses in T cells, but the extent to which response to RVd therapy affects T cells over long periods of sustained treatment may differ.

In summary, we profiled the effects of each RVd drug on T cells using TEA-seq as a discovery platform for drug effects and mechanisms. Each drug induces substantial changes to the transcriptional, epigenetic, and cell surface proteome of T cells. We find that these changes have important implications for T cell function, which could affect patients under monotherapy or combinatorial therapy. Though each drug has a distinct molecular target, we found that there are substantial overlaps and conflicts that arise with co-administration and suggest a possible sequence of administration that could increase the efficacy of RVd therapy. We also show that each drug exerts effects on T cells that could help or hinder post-ASCT, T cell dependent therapies. Further investigation of T cell state in RRMM could build on this work to establish whether the acute observations in this study reflect changes that occur *in vivo* as a result of chronic RVd therapy. Ultimately, both disease and therapy shape the immune landscape. Therapies that rely on a patient’s own cells as critical effectors require an understanding of the molecular effects of drug treatments on key cellular functions. This study demonstrates how timing and sequence of administration may affect the design of new therapies that are consistently effective to drive better outcomes for patients.

In addition to our analysis and observations, we invite readers to pursue their own hypotheses using these datasets and the interactive resources (fig. S11, note S3) available at https://apps.allenimmunology.org/aifi/insights/tcell-vrd/.

## Materials and Methods

### Study Design

The objective of the study was to examine the effects of bortezomib, lenalidomide, and dexamethasone on T cells isolated from healthy adults. The study comprised a cross-sectional analysis of peripheral blood samples of healthy donors. No therapeutic intervention was administered to study participants for the purpose of the study.

### Human PBMC collection

Freshly-drawn whole blood specimens were purchased from sample collections conducted under IRB-approved protocols by Bloodworks Northwest with signed informed consent forms from all donors. Usage of all samples from human subjects are specified in table S1. PBMCs were isolated using Ficoll Premium (GE Healthcare) and stored in Cryostor10 (StemCell Technologies) in liquid nitrogen until use.

### T cell isolation

T cells were enriched from PBMCs by negative selection prior to culture and drug stimulation using RapidSpheres (STEMCELL Technologies). Detailed methods are provided in Supplementary Methods.

### Drug stimulation of isolated T cells

T cells were centrifuged for 5 min at 400g and resuspended in growth medium (RPMI 1640 (Gibco), 20% FBS, 1X Pen/Strep, 50 ng/mL IL-2, 5 ng/mL IL-7, 5 ng/mL IL-15). Drugs were added to cells at the indicated concentrations in Nunc™ 96-well polystyrene round-bottom microwell plates (Thermo Fisher Scientific) in a total volume of 200 µL/well. Plates were incubated at 37°C with 5% CO2 for 4, 24, or 72 hours after drug or control treatment.

### Flow cytometry acquisition and analysis

Data were acquired on a 5 laser Cytek Aurora spectral cytometer using Cytek SpectroFlo software version 2.2.0.4. Intracellular and extracellular staining protocols and panels, as well as compensation and processing details are available in Supplementary Methods. Comparisons of MFI values between conditions from replicates across multiple subjects were performed using paired T-tests with FDR adjustment for multiple hypothesis testing were performed on transformed values. FDR values < 0.05 were considered significant.

### TEA-seq

To accommodate the TEA-seq protocol, thawing and treatment was carried out in a staggered-start reverse time course so that all samples could be collected simultaneously for processing. Samples collected following the viable CD3+ T Cell sort were centrifuged at 400g for 5 min at 4C and resuspended in PBS. Each sample was then individually stained with a panel of 55 oligo-conjugated antibodies (table S15) as previously described (22). Each sample was also stained with a unique Cell Hashing Antibody (table S16) to enable sample pooling and reduce well-to-well effects (60). Following a 30 min incubation, each sample was washed 3 times with PBS + 2% BSA to remove unbound antibodies, counted, and pooled. The hashed cell pool was then tagmented, and single cell GEX, ATAC, and ADT libraries were made as previously described (22). HTO (hash tag-derived oligo) primers were added to the pre-amplification PCR mix, and HTO libraries were prepared from the captured cell hashing antibodies in the same way as the ADT libraries. Final libraries were sequenced on an Illumina NovaSeq 6000.

### DEG analysis

After cell type identification, differential gene expression analysis was performed using the MAST package to compare expression distributions between each drug treatment condition and the DMSO-only condition at each timepoint (4, 24, and 72 hours). Further details are available in the Supplementary Methods.

### Pathway enrichment analysis

We used the fgsea package to perform Gene Set Enrichment Analysis (GSEA) for pathway sets obtained from MSigDB (Hallmark Pathways) and Reactome (Reactome Pathways). Prior to use of fgsea, DEG results from MAST were ranked based on nominal *P*-values and direction of effect, such that the most significantly up-regulated genes were first and most significantly down-regulated genes were last in order. Enrichment tests for Hallmark Pathways and Reactome Pathways were performed separately.

### DEG comparisons between treatment conditions

DEG sets were compared between treatment conditions within each cell type using hypergeometric tests. For each condition, we compared the top 500 up- and down-regulated DEGs. Additional details are available in the Supplementary Methods.

### Differential epitope detection analysis

Differential epitope analysis was performed for each cell type, drug, and time point relative to the DMSO control using normalized ADT count data (Seurat NormalizeData, normalization.method = "CLR", margin = 2). As with DEG analysis, cells were downsampled evenly across timepoints to the minimum cell number by drug and cell type. Epitope expression was modeled linearly as exp ∼ treatment with DMSO as the reference treatment level.

Treatment coefficient *P*-values were adjusted using the Benjamini & Hochberg correction (61).

### DAP analysis

Cells were distributed into separate ArchR projects according to cell type. Peaks were identified using using ArchR’s addGroupCoverages and addReproduciblePeakSet function with parameter groupBy = "Sample". Insertion site counts were tabulated using ArchR’s addPeakMatrix function. Treatment-specific ArchR projects were generated using only cells from each treatment condition at each time point and the relevant timepoint-matched DMSO control cells, then DAPs were identified using ArchR’s getMarkerFeatures function.

### Motif enrichment analysis

For each cell type, peaks were annotated with cisBP motif locations using ArchR’s addMotifAnnotations with parameters motifSet = "cisbp" and the default parameter of version = 2 to use cisBP Version 2 motifs. To identify enriched motifs, DAPs were divided into peaks with increased accessibility (false discovery rate [FDR] < 0.05 and Log2FC > 0) and decreased accessibility (FDR < 0.05 and Log2FC < 0). For DAPs in each direction, hypergeometric tests for enrichment were performed using ArchR’s peakAnnoEnrichment function.

### Statistical Analysis

Statistical analyses were carried out using R v3.6.3 or later. Differentially expressed gene (DEG) tests were performed using the two-part hurdle model implemented in the MAST package including the cellular detection rate as a cofactor, as described in the MAST publication (62). *P*-values were corrected for multiple hypothesis testing using the FDR adjustment (61). FDR values < 0.05 were considered significant. Gene Set Enrichment Analysis was performed using the ranked set correlation approach implemented in the FGSEA package (63). *P*-values were corrected for multiple hypothesis testing using the FDR adjustment. FDR values < 0.1 were considered significant. Direct comparisons of DEG sets (i.e., between drug treatment conditions) were performed using hypergeometric tests with FDR adjustment for multiple hypotheses. FDR values < 0.01 were considered significant. Tests for differentially detected epitopes (DDE) were performed on values normalized using centered log ratio (CLR) transformations as implemented in the Seurat package (24). Normalized values were then compared using linear models in R, with FDR adjustment for multiple hypothesis testing. FDR values < 0.05 were considered significant. Differentially accessible peaks (DAP) were identified using the Wilcoxon tests implemented in the ArchR package (64) with FDR adjustment for multiple hypothesis testing. FDR values < 0.05 were considered significant. Motif enrichment in sets of peaks was performed by comparison of DAP to matched background peak sets using hypergeometric tests with FDR adjustment for multiple hypothesis testing. FDR values < 0.01 were considered significant. Tests for changes in flow cytometry activation marker proportions were performed by CLR transformation of proportional values as implemented in the compositions package for R (65). Paired T-tests with FDR adjustment for multiple hypothesis testing were performed on transformed values. FDR values < 0.05 were considered significant.

## Acknowledgments

The authors thank Melinda Angus-Hill, Claire Gustafson, Emma Kuan, Adam Savage, and Philip Greenberg for reading and providing thoughtful feedback on the manuscript. We also thank Nina Kondza, Nina Estep, Kenny Dang, and Leila Shiraiwa for operational support, Anthony Cicalo and Tao Peng for data processing assistance, and the Allen Institute for Immunology Software Development team for computational support. We utilized generative artificial intelligence (Claude Science) to assist with literature search and summaries. This research was supported by the Allen Institute, founded by Jody Allen – chair and co-founder of Allen Family Philanthropies, and the late Paul G. Allen – investor, philanthropist, and co-founder of Microsoft. We gratefully acknowledge their vision and generosity, which make this work possible.

## Author contributions

Conceptualization: LTG, TRT, PJS

Data Curation: UK

Formal Analysis: LO, LTG, SRZ, ZH

Funding Acquisition: TB

Investigation: WC, JG, CS, MDAW, VH

Methodology: WC, JG, PR, VH

Visualization: LTG, LO

Project administration: JM, MK

Supervision: TRT, PJS, XL, JR, EC, TB

Writing – original draft: LTG

Writing – review & editing: LTG, SBM, TRT, SRZ, GLS, LO, EWN, PJS

## Data and Code Availability

Raw data for TEA-seq experiments is in the process of depositing to dbGaP for controlled access at accession phs003430.v1.p1. Processed TEA-seq data is available on the Gene Expression Omnibus at study accession GSE236422. Code used for data analysis is available on Github at https://github.com/aifimmunology/repro-vrd-tea-seq/.

## Supplementary Materials

### Supplementary Methods

#### Cryopreserved PBMC thawing

Cryopreserved PBMC samples were thawed into pre-warmed AIM V media (37°C) (GIBCO), then resuspended in 5 mL of cold AIM V (4°C) and counted on the Nexcelom Cellaca MX cell counter. Individual samples were resuspended in Dulbecco’s phosphate-buffered saline (DPBS) (Costar) at a concentration of 10 million cells per mL.

#### T cell isolation by negative selection

Thawed PBMCs were resuspended in separation buffer (DPBS containing 2% FBS, Sigma Aldrich, and 1 mM EDTA, ThermoFisher Scientific) at 50 million cells per mL. Samples were transferred to a 5 mL polystyrene round-bottom tube. Isolation Cocktail (STEMCELL Technologies) was added to each sample at 50 µL per mL, mixed, and incubated at room temperature for 5 min. RapidSpheres (STEMCELL Technologies) were vortexed for 30 seconds and 40 µL per mL were added to each sample. A separate buffer was added to samples to fill a total volume of 2.5 mL. After mixing the sample gently by pipetting, the tubes were placed into the EasySep™ magnet (STEMCELL Technologies) and samples were incubated at room temperature for 3 min. Isolated T cells were collected by decanting the enriched cell suspension into a new tube.

#### Extracellular flow cytometry staining

T cells were washed and resuspended in 200 µL FACS wash buffer (2% BSA and 0.5 mM EDTA in DPBS) and transferred to a new 96-well round-bottom plate, then centrifuged at 450g for 5 min, and cell pellets were resuspended to a final volume of 100 µL with a viability mix containing 1:400 FVS510 viability dye (BD, 1 µg/mL) in FACS wash buffer and incubated for 30 min at 4°C protected from light. After an additional wash with 200 µL FACS wash buffer, samples were resuspended to a final volume of 100 µL with extracellular antibodies in FACS wash buffer (table S12). Samples were then mixed and incubated for 30 min at 4°C, protected from light prior to data acquisition.

#### Intracellular flow cytometry staining

For experiments using intracellular staining (including Ikaros and Aiolos), cells labeled by extracellular staining described above underwent two additional washes using 200 µL FACS wash buffer, then were resuspended in 200 µL Foxp3 Fixation/Permeabilization working solution (BioLegend) for 30 min at room temperature, protected from light. Samples were centrifuged at 750g for 5 min at room temperature. This was repeated twice by discarding the supernatant and resuspending the pellets in 200 µL 1X permeabilization buffer and centrifuging at 400g for 5 min at room temperature. Intracellular antibodies (table S13) were added in 100 µL total volume, and cells were incubated for 30 min at room temperature, protected from light. Cells were then washed twice in 200 µL 1X permeabilization buffer, then were resuspended in 200 µL FACS wash buffer prior to data acquisition.

#### Flow cytometry compensation and processing

Compensation matrix generation and spectral unmixing was completed by creating single color controls using healthy resting PBMCs (table S14). Single color controls were stained with the same protocol. Spectral unmixing was calculated in the SpectroFlo instrument software (Cytek, Version 2.1.1). Flow Cytometry Standard (FCS) files from the Cytek Aurora spectral cytometer were exported in FCS 3.1 format. FCS files were manually gated to remove doublets, debris, and dead cells using FlowJo (BD Biosciences, Version 10.8.1).

#### TEA-seq data processing

After sequencing, scRNA-seq and scATAC-seq libraries were aligned, and transcripts were counted using CellRanger Multi to generate transcript count matrices and unique fragment alignments for each cell barcode. ADT and HTO count matrices were generated using BarCounter v1.0 (69). RNA, HTO, and ADT count matrices were combined and demultiplexed for each treatment condition using HTO count distributions with BarMixer (69). Additional scripts utilizing BedTools, awk, and ArchR were used to demultiplex scATAC-seq fragments for each sample and perform initial quality control analysis. Cell barcodes were filtered to select cells with the following characteristics: > 500 scRNA-seq genes per cell and > 1,000 unique scATAC-seq fragments per cell. Initial analysis of the scRNA-seq data was performed using Seurat to cluster cells and identify non-T cell contamination and doublets. In total, 181,518 cells were considered, 37,690 cells (20.8%) were excluded based on QC criteria and clustering, and 143,828 cells (79.2%) were retained for downstream analysis.

#### Cell-type labeling

Normalized ADT count distributions for CD4 and CD8a antibodies were inspected from each sample to identify cutoffs for CD4-positive and CD8-positive populations. Cells were then separated into CD4 and CD8 T cell classes for downstream labeling steps. For each class, scRNA-seq data was analyzed using Seurat. Counts were transformed using SCTransform, and label transfer was performed using the PBMC cell type reference (Azimuth). To constrain label transfer, reference cells were filtered for those with a top-level cell type label of either CD4 T or CD8 T. Labels were transferred using Seurat’s FindTransferAnchors and MapQuery functions for CD4 and CD8 cells and respective references. After label transfer, all CD4 or CD8 cells were integrated across samples using Seurat functions SelectIntegrationFeatures, PrepSCTIntegration, FindIntegrationAnchors, and IntegrateData. Dimensionality reduction was then performed using RunPCA and RunUMAP, and Leiden clustering was performed using FindNeighbors and FindClusters. Marker gene expression and transfer-based labels (above) were used to assign cell types to each of the CD4 and CD8 Leiden clusters.

#### MAST DEG analysis

For each cell type, we downsampled cell across all treatment conditions based on the lowest number of cells in any condition (additional considerations to this approach are discussed further in note S2). To be considered for differential expression testing, a gene must have been detected in at least 10% of cells in either the drug or DMSO control to be considered for analysis. Then, using the downsampled cells and genes that passed detection cutoffs, we tested for differential expression using MAST using the following formula:

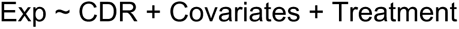

Which included inclusion of Cellular Detection Rate (CDR) to be able to jointly estimate nuisance factors and treatment effects.

#### Gene sets for comparative analyses

For comparison to TEA-seq DEGs induced by dexamethasone treatment, we curated differential gene expression sets from previous literature and databases related to dexamethasone treatment (70–76) and the Gene Ontology (77,78).

For use in GSEA analyses, Reactome Pathways were preprocessed to retain pathways that were within 3 levels of nesting within the Reactome pathway ontology from top-level pathway groups (29 groups, e.g., Autophagy, Metabolism of proteins, and Signal Transduction), resulting in 1,350 total Reactome gene sets used for analysis.

#### DEG comparisons by hypergeometric tests

To account for differences in the number of cells in different conditions, we compared the top 500 up- and down-regulated DEGs. The number of overlapping genes between each set of DEGs was counted and identified the number of common genes that were tested in both conditions. We then used the R function phyper to calculate the probability of overlap between gene sets using the parameters q = N overlapping genes - 1, m = 500 (number of genes in Set A), n = number of common genes not in Set A, k = 500 (number of genes in Set B), and lower.tail = FALSE. After computing all tests, *P*-values were adjusted for multiple hypothesis testing using the p.adjust function in R with the parameter method = "BH" for the Benjamini and Hochberg method.

#### Data analysis and visualization tools

Data analyses were performed using R v3.6.3 or later (79). For general data handling, we utilized the following packages: data.table v1.14.6 (80), dplyr v1.0.10 (81), purrr v1.0.0 (82), tibble v3.1.8 (83), tidyverse v1.3.2 (84), and tidyr v1.2.1 (85). For single-cell genomics analyses, we utilized ArchR v1.0.2 (64), clustree v0.5.0 (86), fgsea v1.24.0 (63), MAST v1.24.0 (62), Seurat v4.3.0 (24), and the readGMT function from GSEABase v1.60.0 (87). For data visualization, we utilized cowplot v1.1.1 (88), ggplot2 v3.4.0 (89), ggrastr v1.0.1 (90), ggrepel v0.9.2 (91), and Adobe Illustrator v27.8.1. For flow cytometry analysis, we used FlowJo v10.9.0.

### Supplementary Notes

#### Supplementary Note 1. Dose selection based on pharmacokinetic reports

To select appropriate drug treatments to induce T cell state perturbation, we examined pharmacokinetic reports of each individual drug administered at doses used in Multiple Myeloma treatment. Bortezomib administered subcutaneously at 1.3 mg/m2 had a steep decline from a maximum plasma concentration of 54.04 nM (92), resulting in a median AUClast of 403 nM h over the course of 72 hours. In culture, where there is no liver clearance of bortezomib over the course of 72 hours, 5.6 nM of exposure over the course of 72 hours would provide the same AUC exposure over time, assuming no loss of bortezomib during the time course.

Lenalidomide administered orally at 25 mg has a maximum plasma concentration of 2,190 nM (93), with less rapid initial clearance than that observed with bortezomib. Dexamethasone administered orally at 20 mg had a maximum plasma concentration of 694.5 nM (94). High concentrations of Bortezomib (10 nM) caused complete loss of T cells in culture over 24 hours in initial in vitro testing (data not shown), so titrations of 5 nM, 2.5 nM, and 1 nM, close to the dose encountered over 72 hours, were tested for use in time course experiments; Lenalidomide was tested at 1500 nM, 1000 nM, and 500 nM; and Dexamethasone was tested at 100 nM, 50 nM, and 25 nM concentrations.

#### Supplementary Note 2. Cell type resolution and differential expression testing

The differential gene expression analysis carried out in this study relied on the MAST method, which allows for use of gene detection and models both the discrete and continuous aspects of single-cell gene expression data. While MAST is a powerful tool for identifying differentially-expressed genes, we have noticed that the number of statistically significant results is correlated with the number of cells used as input for these tests. Thus, comparisons between tests with different input cells can be misleading. However, we would like to maintain as much power as we can to describe the effect of RVd therapy drugs. To balance resolution and power, we present our cell types at a resolution that allows for robust differential expression tests. We are able to identify finer cell types in our dataset (e.g. we can separate CD8 Memory into CD8 Effector Memory, CD8 Central Memory, and CD8 TEMRA cells), but doing so reduces the number of cells in our comparisons below a useful threshold for detecting differences in gene expression. Compounding the issue of cell type abundance is the difference in number of cells per treatment condition that were obtained in our experiments. To capture the effects of our cell culture conditions, we always compare our drug treatment samples to the appropriate DMSO-only control sample (e.g., Bortezomib at 72 hr to DMSO at 72 hr; Lenalidomide at 4 hr to DMSO at 4 hr). We have found that having a balanced number of cells is helpful for obtaining good MAST differential expression results. Thus, we also consider downsampling to match the number of cells compared in the foreground (treatment) and background (control). One way to balance comparisons across cell type, time, and treatments is to downsample all comparisons to the lowest number of cells available in any single condition. However, this leaves us with very few cells, and results in low power. The compromise we have arrived at is to downsample with each cell type at a moderate resolution (as described above) within each drug and timepoint.

This allows us to compare the effect of each drug across time points within each cell type - making it apparent, for example, that the effect of bortezomib increases over time within some cell types, or that the maximum effect of lenalidomide occurs at 24 hours (of the timepoints tested). However, readers should be cautioned that comparisons *between* drugs or *between* cell types are not strongly supported by this strategy.

#### Supplementary Note 3. Interactive Visualization Tools

We provide a suite of cloud-based interactive visualization application to facilitate exploration of the RVd drug perturbation T-cell TEA-seq results at https://apps.allenimmunology.org/aifi/insights/tcell-vrd/.

Here we describe the app’s interactive features.

#### DEG Explorer

The DEG Explorer offers a comprehensive look at drug-perturbed genes both within and across cell types and individual drug treatments (fig. S1A). The design is intended to allow users to see the overall scale of impacted genes per drug (volcano plots) and help users explore biological relevance of changes through flexible gene and gene set annotation and by providing gene summary and ontology information. For example, for a given drug and celltype, users can overlay labels for genes in a Hallmark Pathway to see if there are coordinated changes related to a known biological process. Users can alternatively overlay their own set of annotations to explore their own hypotheses or compare findings from other studies. By selecting a specific gene, users can view differential expression across all celltypes and drug treatments in heatmap form. This view can afford insights into which drug effects may be cell-type specific as well as show how genes are affected differently by drugs, where concordant vs discordant regulation would have different implications in the context of combination therapy.

Usage: A volcano plot panel displays differential gene expression results for a selected drug, timepoint, and cell type relative to the DMSO control for an overview of transcriptional changes. Clicking a gene on the plot (or choosing from the provided dropdown) will update a heatmap panel summarizing the gene’s DEG results for all cell types and drug treatments. Gene selection will also populate the reference panel containing a gene summary and ontological details (Source: http://mygene.info).

Genes of interest may be annotated on the volcano plot by expanding the annotation panel and choosing various options (fig. S1B). Individual genes or entire Hallmark Genesets can be selected for annotation via dropdown. Users can provide their own gene list via copy/paste into the provided form. Genes can also be chosen directly from the volcano plot by clicking- or box-selection, e.g., to bulk label the top differential results. Annotations will remain manually cleared, allowing users to to toggle the underlying data for a particular set of features.

**Supplementary Figure 1:**
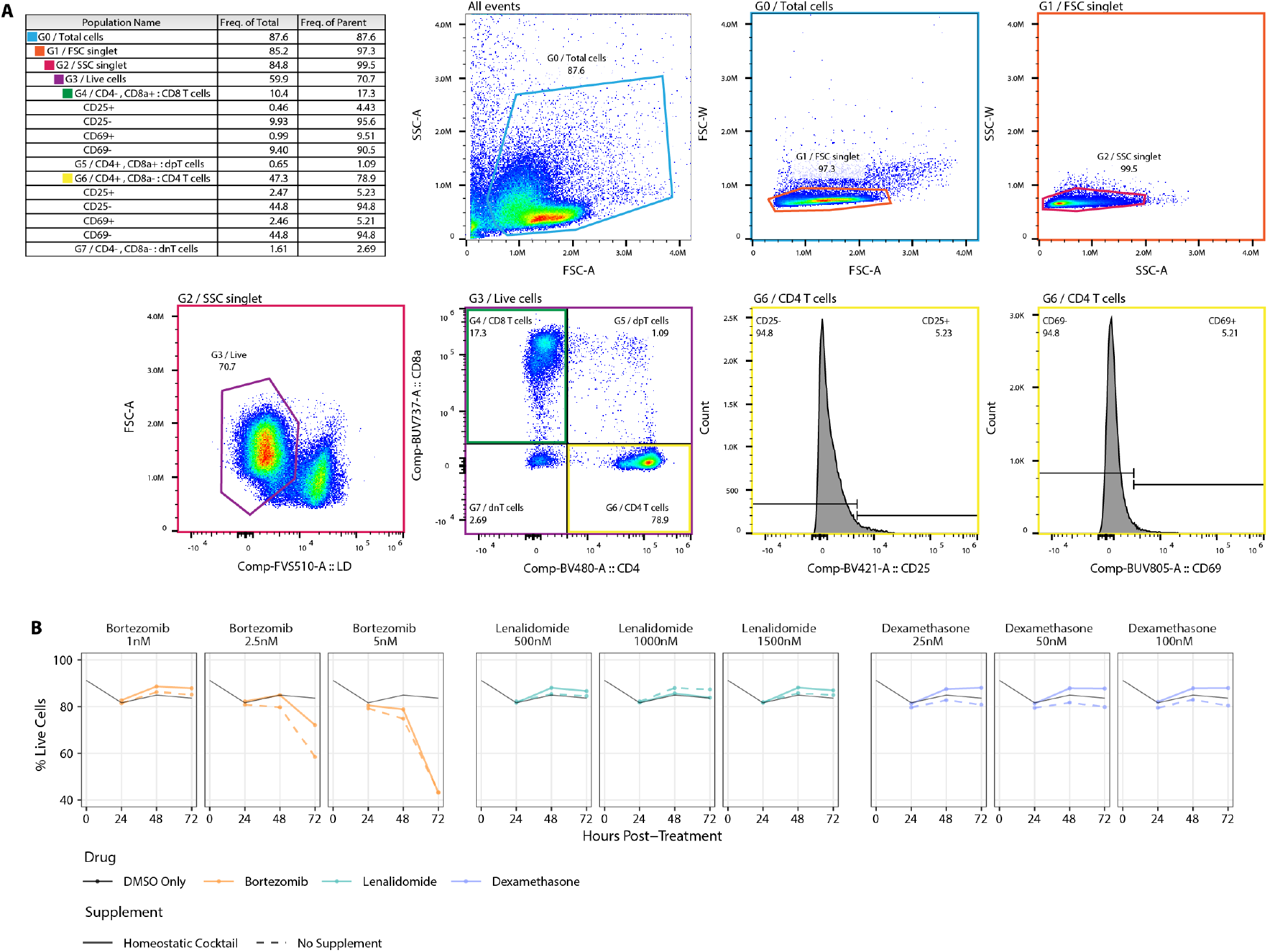
Cell survival under drug treatment conditions. **A.** Flow cytometry gating scheme and example gating diagrams for quantification of live cells and selection of CD4 and CD8 T cells to assess activation marker expression (CD25 and CD69). **B.** Line plots demonstrating cell survival over the course of 72 hours under increasing drug treatment concentrations. In each panel, a single drug concentration (colored lines) is compared to DMSO only controls. Note that the DMSO control values are the same across all panels. Cells treated with the homeostatic cytokine cocktail (IL-2, IL-7, and IL-15) are plotted using solid lines. Non-supplemented cells are plotted using dashed

**Supplementary Figure 2:**
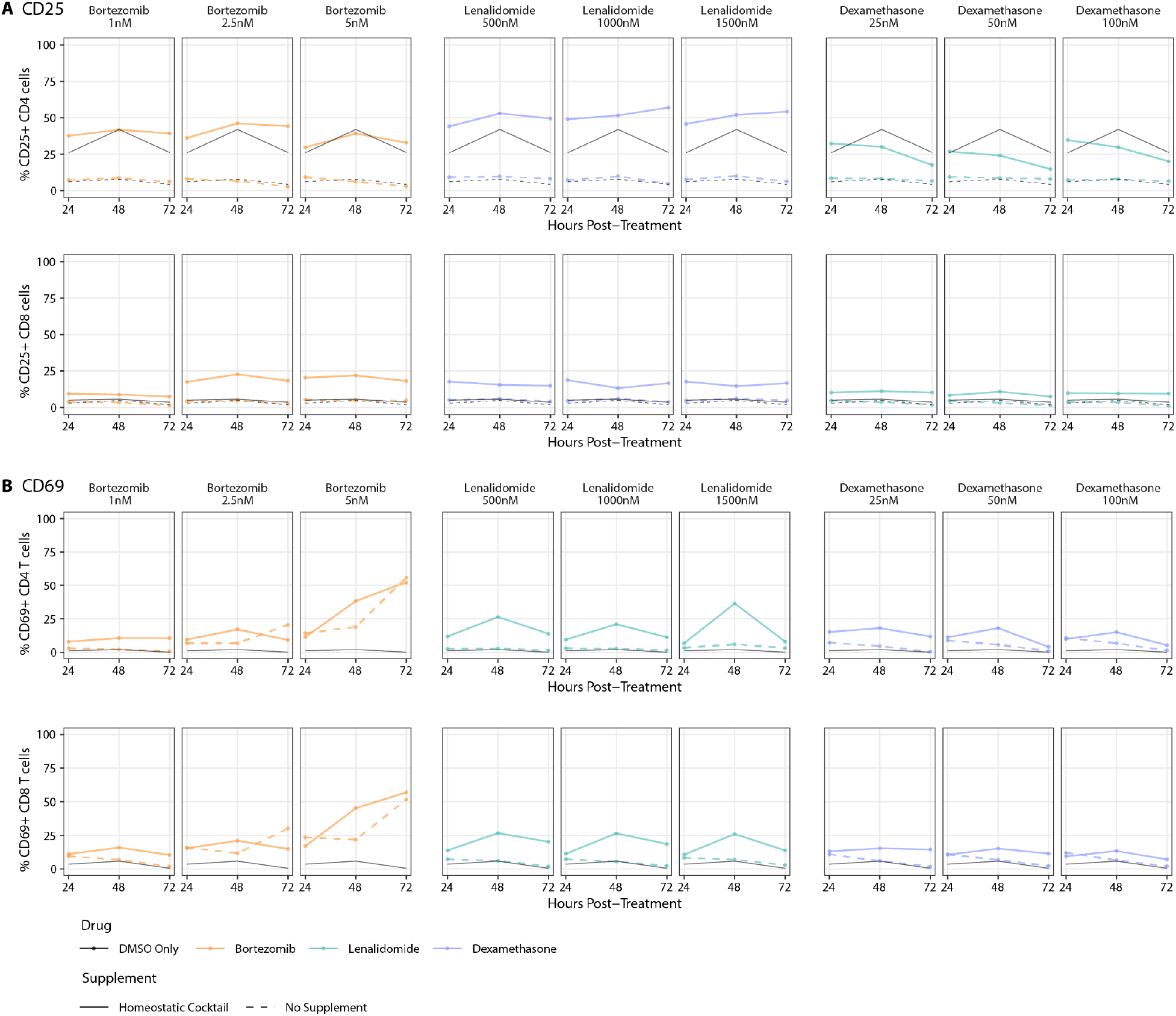
Cell activation under homeostatic cocktail and drug treatment conditions. **A.** Line plots showing the fraction of CD4+ T cells (top row) or CD8+ T cells (bottom row) that showed expression of the T cell activation marker CD25 over the course of 72 hours. In each panel, a single drug concentration (colored lines) is compared to DMSO only controls. Note that the DMSO control values are the same across all panels. **B.** As in (**A**) for expression of the T cell activation marker CD69.

**Supplementary Figure 3:**
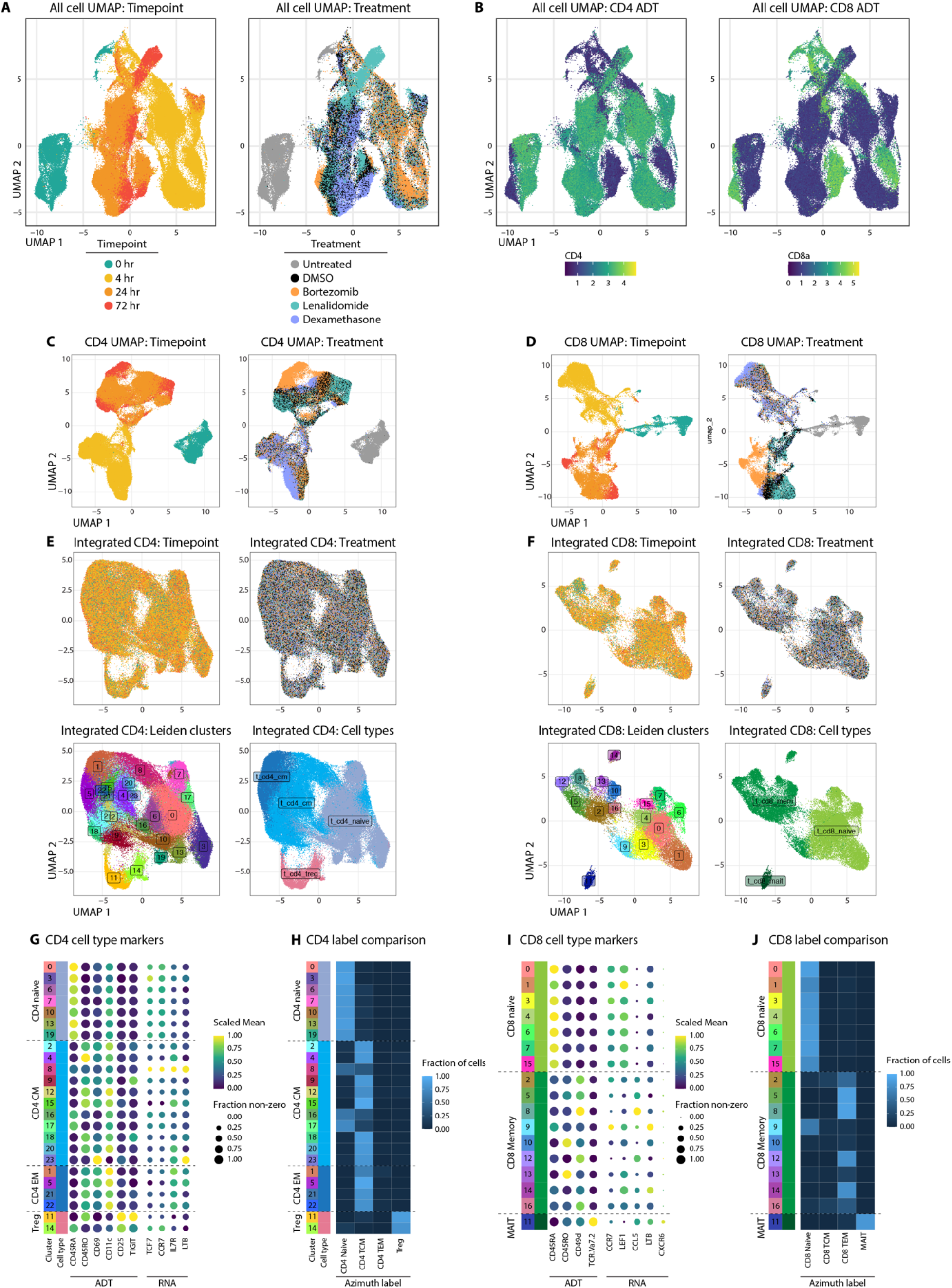
TEA-seq Dataset Overview and cell type labeling. **A.** UMAP plots all available cells after QC filtering and removal of non-T cells as described in Methods (n = 143,828 cells). UMAP coordinates were generated using 3-way Weighted Nearest Neighbors embedding, as described in (24). Cells are colored according to the timepoint (left) or drug treatment (right) for each sample. **B.** UMAP embeddings, as in (**A**), colored based on CLR-transformed CD4 (left) or CD8 (right) antibody-derived tags to delineate CD4 and CD8 T cells. **C.** UMAP projections for all CD4+ cells using scRNA-seq data alone with default settings and colored based on timepoint (left) or drug treatment (right). **D.** as in (**C**), but for all CD8+ cells. **E.** UMAP projections for all CD4+ cells using scRNA-seq data alone after performing integration of samples. This causes mixing of all timepoints and treatments and allows unified cell type clustering and identification to be performed across conditions. Cells are colored based on timepoint (top left), drug treatment (top right), Leiden clustering results (bottom left), or final cell type labels (bottom right). **F.** as in (**E**), but for all CD8+ cells. **G.** A dot plot displaying selected marker genes (columns) for CD4+ cell types in each of the Leiden clusters labeled in (**E**) (rows). Both ADT-derived and RNA-derived expression values are displayed (indicated at bottom). Dots are colored based on scaled mean expression values (mean for clusters divided by the highest mean across all clusters) and are sized based on the fraction of cells in each cluster that have non-zero expression values. **H.** Heatmap displaying the fraction of CD4+ cells in each Leiden cluster (rows) that were labeled with each Level 2 cell type from the Azimuth reference (columns) based on Seurat label transfer. **I.** as in (**G**) for CD8+ cell types. **J.** as in (**H**) for CD8+ cell types.

**Supplementary Figure 4.**
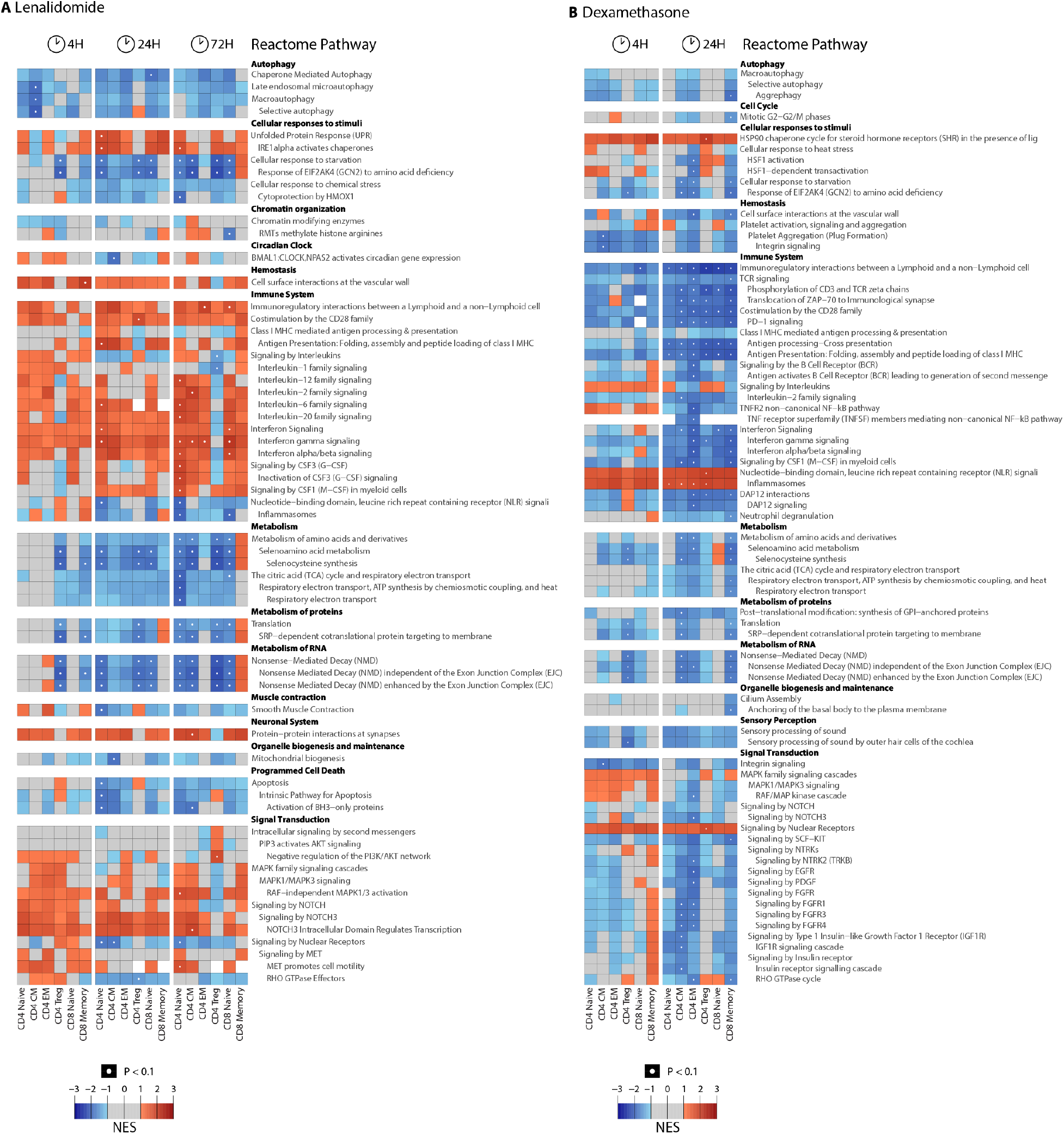
Lenalidomide and dexamethasone Reactome Pathways. Heatmaps showing normalized enrichment scores for Reactome pathways, as in fig. S4, but for comparisons between lenalidomide (**A**) or dexamethasone (**B**) and DMSO treatment at each time point.

**Supplementary Figure 5.**
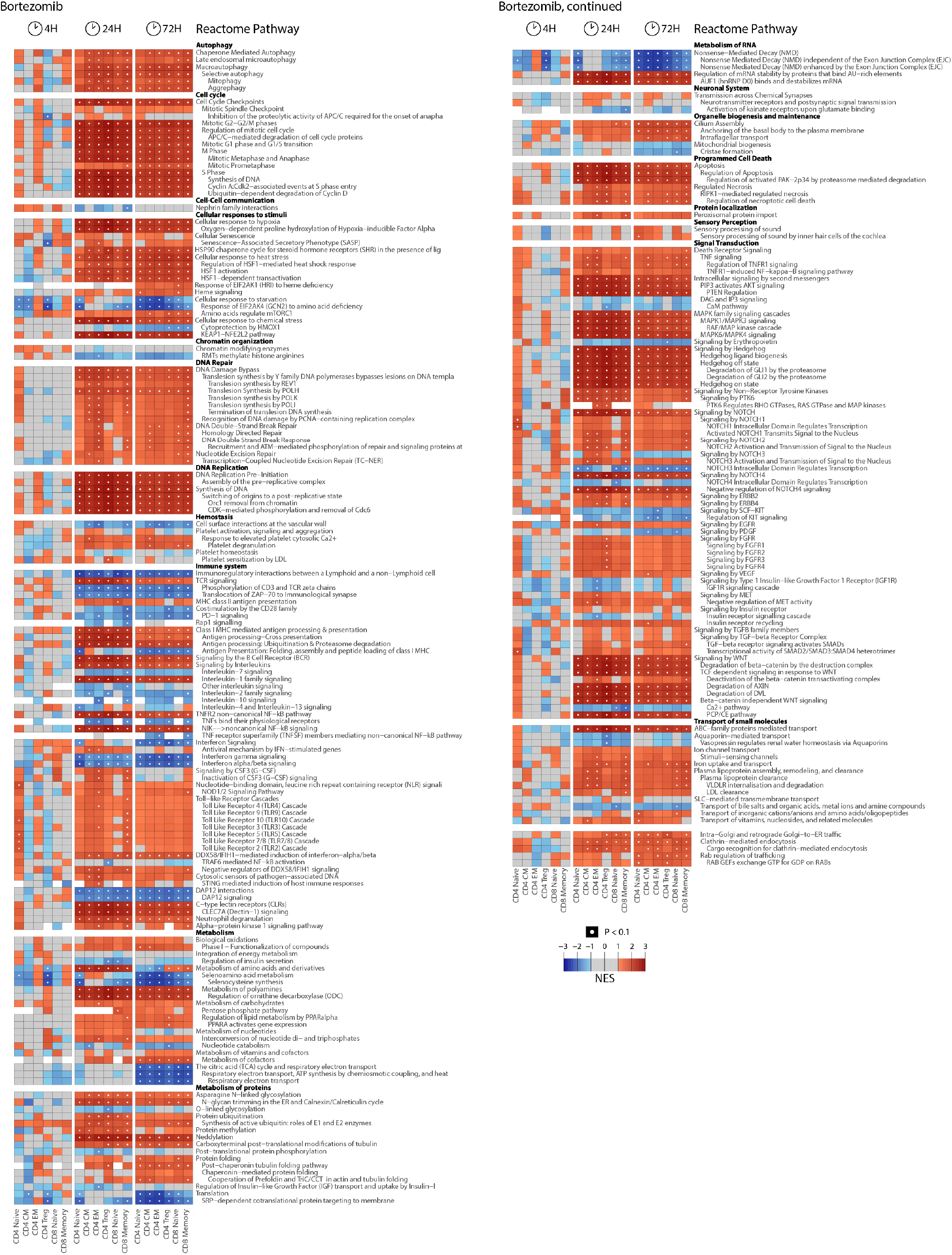
Bortezomib Reactome Pathways. Heatmaps showing normalized enrichment scores (NES) for Reactome pathways (rows) for each cell type (columns) in response to bortezomib at each time point (column groups separated by white space). Reactome pathways are grouped based on their ontological relationships. Labels for sub-pathways are identified relative to parent pathways. NES values > 1 (enriched among up-regulated genes) are represented by red colors, NES values < -1 are represented by blue colors (enriched among down-regulated genes). NES values between -1 and 1 are represented by gray. Enrichment results with an adjusted *P*-value < 0.1 are indicated with a white point. Full results, including exact *P*-values, are provided in table S6.

**Supplementary Figure 6:**
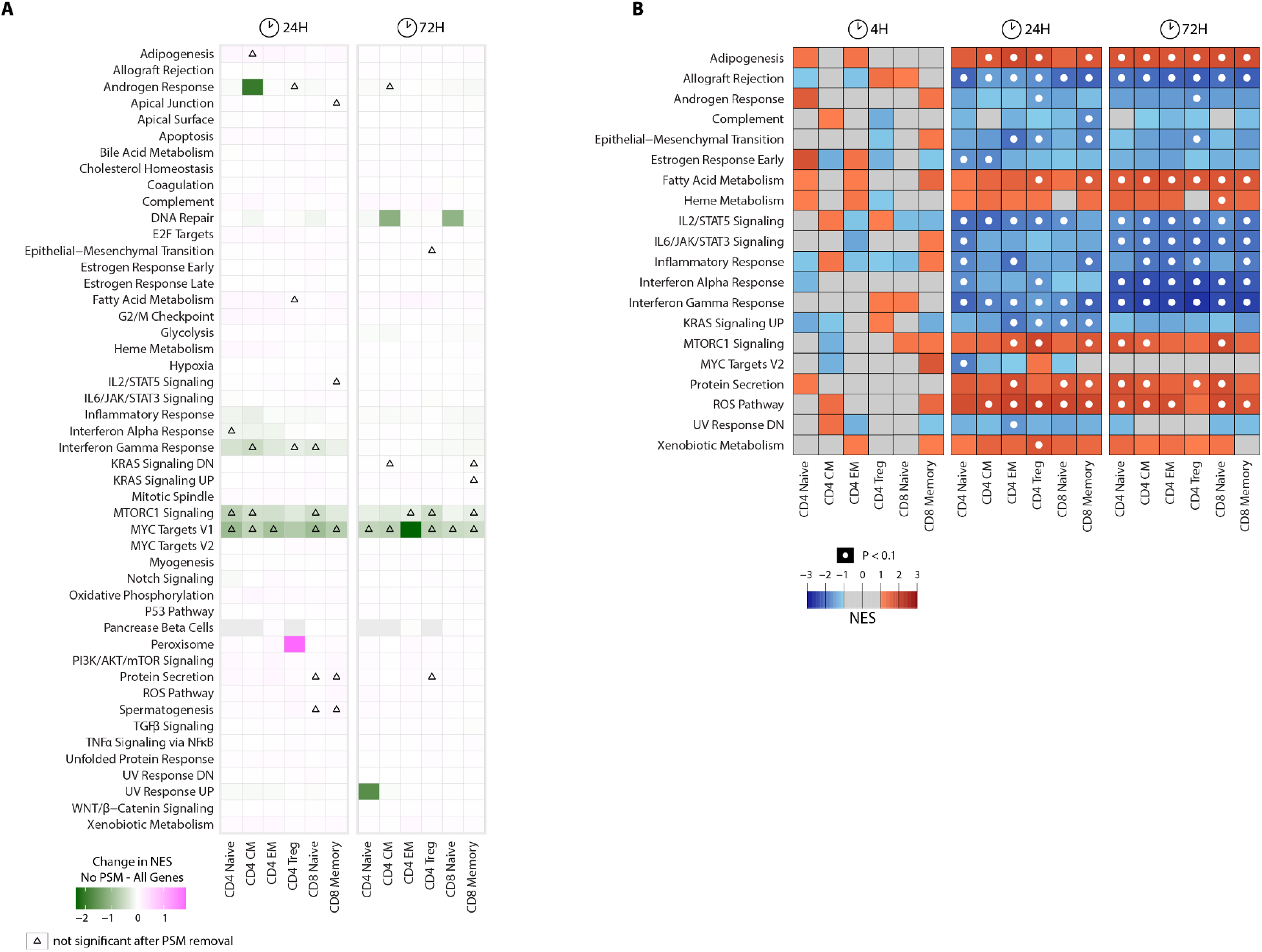
Bortezomib responses without proteasomal genes. **A.** Heatmap showing the difference in NES values among DEGs when proteasomal genes are removed relative to when all genes are present for each Hallmark Pathway (rows) in each cell type (columns). Triangles indicate values that were significant when all genes were included, but lost significance after removal of proteasomal genes. **B.** Heatmap displaying NES values for the same pathways presented in Fig. 2C after removal of proteasomal genes. 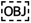

**Supplementary Figure 7:**
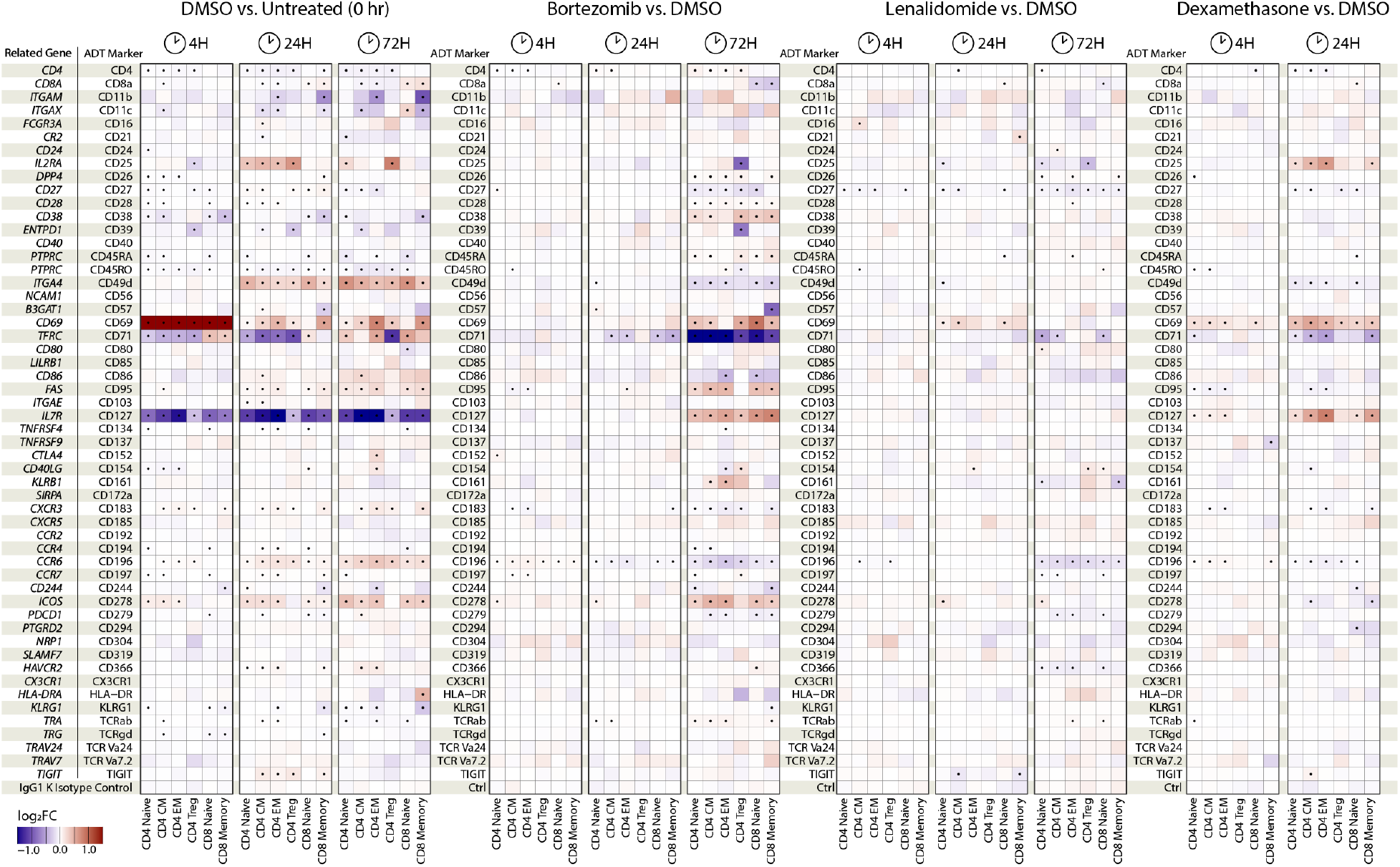
Differential detection of cell surface epitopes in TEA-seq data. Heatmaps displaying the log_2_(Fold Change) in mean surface epitope expression as measured by Antibody-derived tag (ADT) detection on single cells. Each row represents a single antibody marker conjugated to a barcoded oligonucleotide (54 markers, 1 IgG1 K Isotype negative control). Columns represent differential expression in each cell type (labeled at bottom) at the time of point specific (labeled at top). In the first set of panels, DMSO vs. Untreated (0 hr), DMSO control samples from the 4 hr, 24 hr, or 72 hr time point were each compared to the Untreated control sample, which was processed immediately after cell thawing. In all other panels, the drug treatments were compared to the DMSO controls at matched time points (e.g., Bortezomib at 24 hours compared to DMSO at 24 hours). Statistically significant changes are marked with a black point (adjusted P-value < 0.01).

**Supplementary Figure 8:**
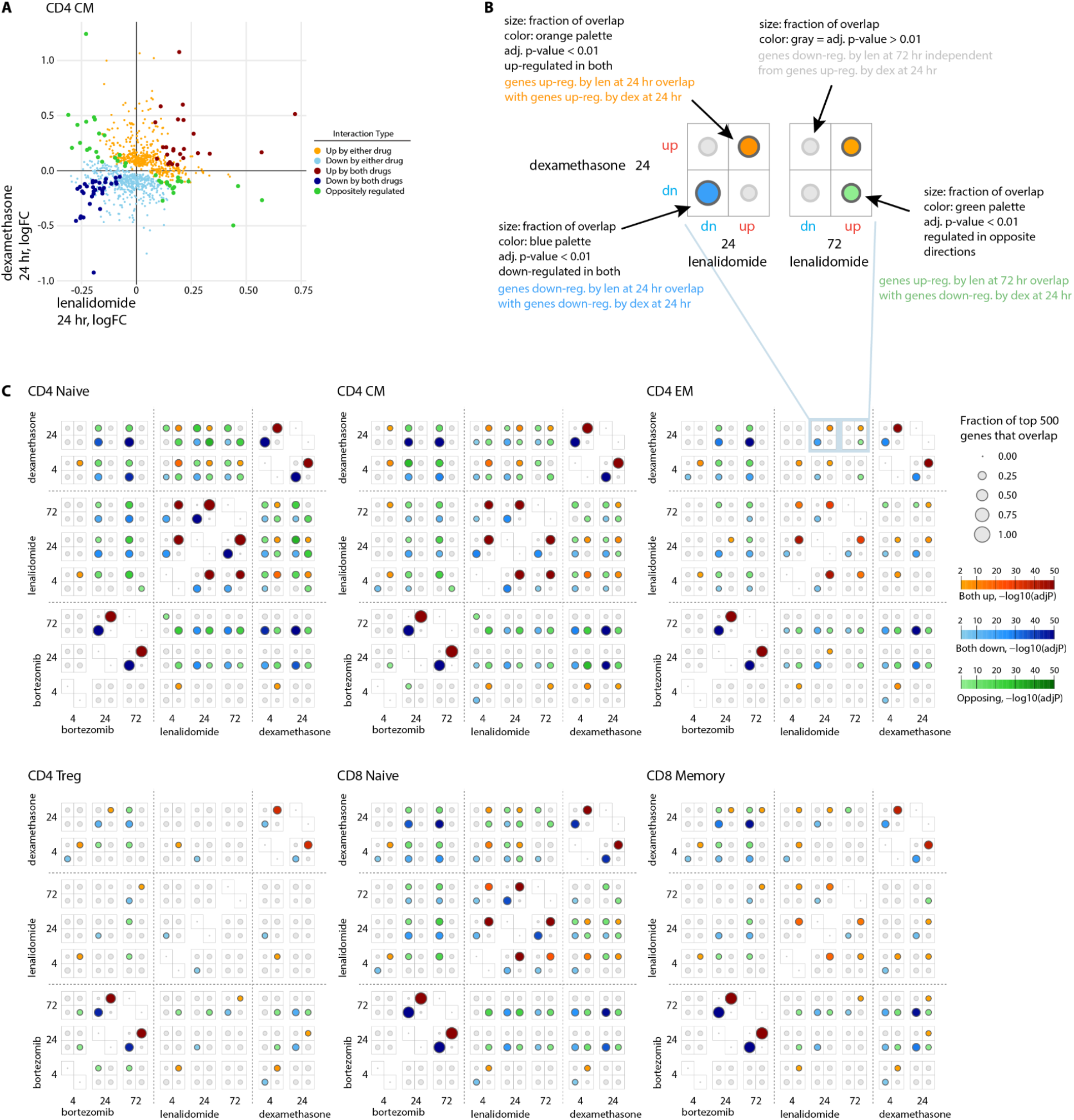
Comparison of DEGs between drug treatment conditions. **A.** Pairwise comparison of log fold change values in CD4 CM cells at 24 hours induced by lenalidomide (x-axis) and dexamethasone (y-axis). Each point represents a single gene. Points are colored based on agreement or disagreement between the two drug conditions: orange, significantly up-regulated by only one of the two drugs; light blue, down-regulated by only one of the two drugs; dark red, up-regulated by both drugs; dark blue, down-regulated by both drugs; green, oppositely regulated (significantly up by one and down by the other). **B.** Plot guide for interpretation of panels in (**C**). **C.** Dot plot displaying the fraction of overlapping DEG between drug treatment conditions. Size of points indicate the fraction of the top 500 genes up- or down-regulated in each treatment condition. Points are colored based on the type of overlap: orange, up-regulation by both conditions; blue, down-regulation by both conditions; green, regulated in opposite directions. 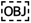

**Supplementary Figure 9:**
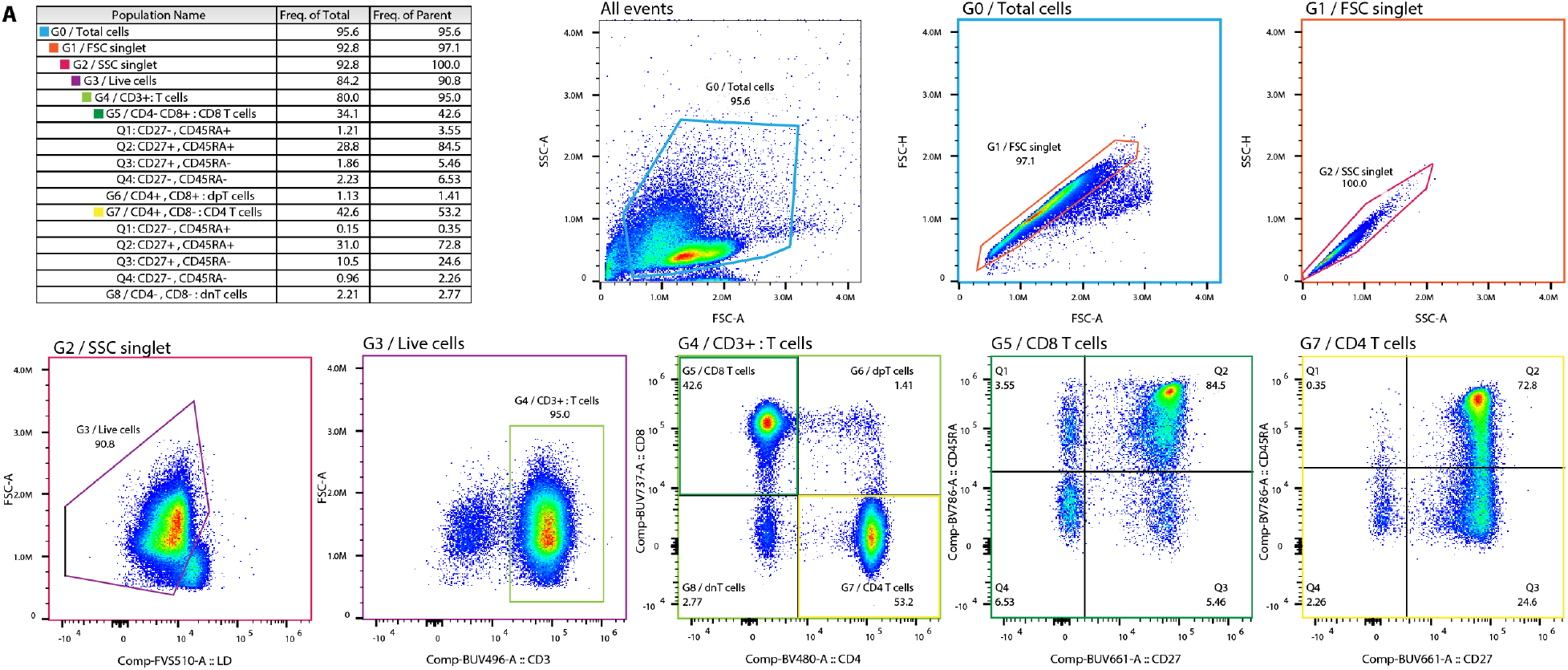
Cell surface marker expression after treatment with each combination of drugs. Example gating diagrams for flow cytometry used to identify CD4 and CD8 T cells for comparison of cell surface marker expression presented in Fig. 6D.

**Supplementary Figure 10:**
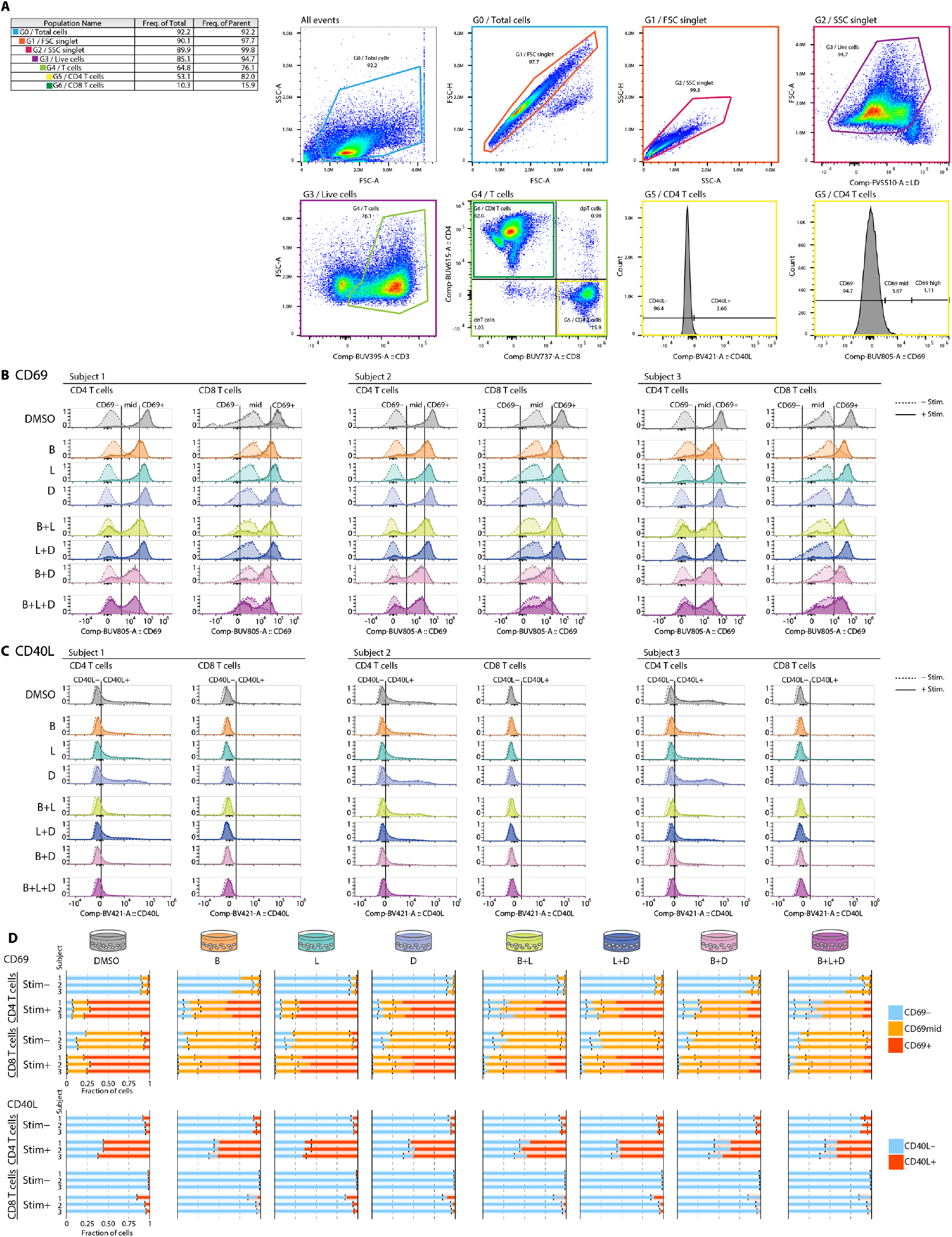
Changes to T cell activation following treatment with each combination of drugs. **A.** Example gating diagrams for flow cytometry used to measure CD40L and CD69 expression on CD4 and CD8 T cell populations. **B.** Histograms showing the CD69 expression in CD4 (left) and CD8 (right) T cells for each subject, grouped by headers at the top of each panel. The CD69 signal derived from unstimulated T cells is shown using a dashed line, while the signal from CD3/CD28 stimulated T cells is shown using a solid line. Each row represents a single treatment condition, specified at the left side of the row: DMSO, DMSO only, B, bortezomib, L, lenalidomide, D, dexamethasone. Vertical lines indicate cutoffs for CD69-negative (CD69-), CD69-mid, or CD69-high (CD69+). For the purposes of statistical tests presented in Fig. 6F, only the fraction of cells in the CD69+ group are considered. **C.** as in (**B**), but for CD40L expression. **D.** Bar plots representing fractional quantification of the cell populations shown in histograms in (**B-C**). Each row represents one of the 3 subjects, and conditions and cell types are grouped as indicated at the left side of the plot. Each column of plots represents a single treatment condition, as indicated above the first row of bar plots. Black vertical lines indicate the fractional values from the DMSO-only control to aid in comparison results to the control condition.

**Supplementary Figure 11.**
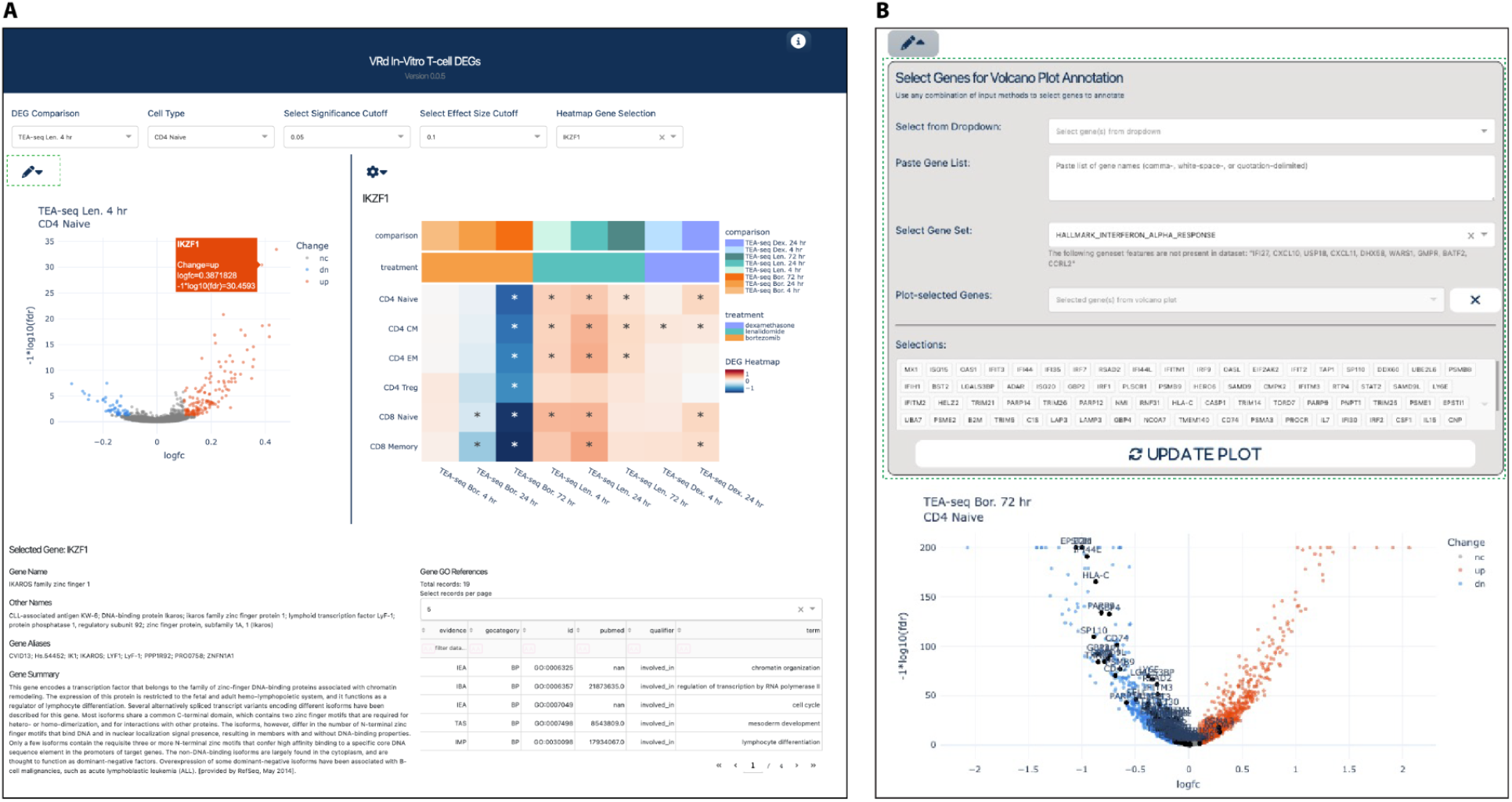
Interactive visualization tools. **A.** An overview of DEG Explorer App interface. The top row of user inputs defines the data selection and plotting options. The left panel volcano plot shows all DEG results for selected drug comparison and cell type. Hover on a point to see the tooltip info, shown for IKZF1 in this example. On gene selection, summarized DEG results for all cell types and drug conditions are plotted in the heatmap panel on the right. The bottom panel gives gene summary information. **B.** Dropdown menu for volcano plot gene annotations showing options for making gene selections.

### Supplementary Tables

**Supplementary Table 1. Human subjects and samples.** List of all human subjects and samples presented in experiments in this study. Column descriptions:

Experiment: Experimental method applied to sample
Figure(s): Figures displaying results derived from sample
Table(s): Tables displaying results derived from sample
Sample ID: Identifier corresponding to original PBMC sample
Sample Source: Organization from which sample was acquired
Subject ID: Identifier for human subject from which sample was acquired Subject Age: Age of subject at time of sample acquisition
Subject Sex: Biological sex of subject
Subject Race: Self-reported race of subject
Subject Ethnicity: Self-reported ethnicity of subject

**Supplementary Table 2. TEA-seq cell metadata and labels.** Metadata associated with each individual T cell that passed quality control threshholds. Column descriptions:

treatment: Drug treatment condition
timepoint: Drug treatment timepoint (in hours)
epi_gate_class: Epitope-based cell class gate (CD4 or CD8)
aifi_cell_type: Cell type labels assigned based on clustering and marker expression
barcodes: Unique universal identifier barcodes for each cell
riginal_barcodes: Original sequence-based barcodes for each cell
well_id: 10x Genomics Chromium well number, in the format: Experiment-PoolChipWell
txn_n_genes: Number of genes detected in transcription data
txn_n_umis: Number of unique molecular identifiers (UMIs) in transcription data
txn_n_reads: Number of reads in transcription data
txn_n_mito_umis: Number of UMIs aligning to mitochondrial genes in transcription data
txn_frac_mito_umis: Fraction of total UMIs aligning to mitochondrial genes in transcription data
epi_n_adt_umis: Number of UMIs from epitope data (antibody-derived tags, ADT)
acc_n_unique: Number of unique fragments in accessibility data
(scATAC)
acc_n_fragments: Number of total fragments in accessibility data
acc_n_mito_frags: Number of fragments aligned to mitochondrial contigs (chrM) in accessibility
data acc_frac_mito_frags: Fraction of fragments aligned to mitochondrial contigs (chrM) in accessibility
data acc_frac_tss: Fraction of fragments aligned to TSS regions in accessibility data (reported by ArchR)

**Supplementary Table 3. Differentially expressed genes.** MAST DEG results for each gene tested in each drug treatment condition compared to the timepoint-matched DMSO control. Column descriptions:

aifi_cell_type: Cell type label
timepoint: Time point of sample collection
fg: Foreground treatment condition
bg: Background treatment condition
n_downsample: number of cells used for downsampling prior to analysis
gene: Gene symbol
nomP: Nominal P-value reported by MAST
logFC: Log fold change reported by MAST
coef_d: Discrete model coefficient reported by MAST
coef_c: Continuous model coefficient reported by MAST
adjP: Adjusted P-value after multiple hypothesis testing correction fg_frac_nz: Fraction of cells in foreground set with non-zero expression of gene
bg_frac_nz: Fraction of cells in background set with non-zero expression of gene
fg_mean_nz: Mean of normalized, non-zero expression values from the foreground set
bg_mean: Mean of normalized, non-zero expression values from the background
direction: Direction of change in expression in foreground relative to background

**Supplementary Table 4. Differentially detected epitopes.** DDE results computed from linear modeling of CLR-transformed ADT count data. Column descriptions:

aifi_cell_type: Cell type label
timepoint: Time point of sample collection
fg: Foreground treatment condition
bg: Background treatment condition
n_downsample: number of cells used for downsampling prior to analysis
feature: Name of the protein epitope measured
estimate: Estimate coefficient from linear model
std_error: Standard Error coefficient from linear model
t_value: T value coefficient form linear model
nomP: Nominal P-value reported by linear model test
fg_mean: Mean of normalized expression values from the foreground set for the feature
bg_mean: Mean of normalized expression values from the background set for the feature
fc: Fold change in expression (fg_mean / bg_mean)
adjP: Adjusted P-value after multiple hypothesis testing correction

**Supplementary Table 5. Differentially accessible peaks.** DAP results computed using wilcoxon tests implemented in ArchR. Column descriptions:

aifi_cell_type: Cell type label
timepoint: Time point of sample collection
fg: Foreground treatment condition
bg: Background treatment condition
chr: Peak region chromosome (GRCh38/hg38 coordinates)
start: Peak region start position (GRCh38/hg38 coordinates)
end: Peak region end position (GRCh38/hg38 coordinates)
Log2FC: Log2 fold change in accessibility (reported by ArchR)
FDR: False Discovery Rate (adjusted *P*-value, reported by ArchR)
MeanDiff: Mean Difference in accessibility (reported by ArchR)

**Supplementary Table 6. Hallmark Gene Set Enrichment Analysis results.** GSEA results for all Hallmark Pathway gene sets tested for enrichment against significant DEGs for each condition. Column descriptions:

aifi_cell_type: Cell type label
timepoint: Time point of sample collection
fg: Foreground treatment condition
bg: Background treatment condition
pathway: Hallmark Pathway name
n_genes: number of genes in Hallmark pathway gene set
nomP: Nominal P-value reported by fastgsea
adjP: Adjusted P-value after multiple hypothesis testing correction
NES: Normalized Enrichment Score reported by fastgsea leadingEdge: Semicolon-separated list of Leading Edge Genes reported by fastgsea

**Supplementary Table 7. Reactome Gene Set Enrichment Analysis Results.** GSEA results for all Reactome Pathway gene sets tested for enrichment against significant DEGs for each condition. Column descriptions:

aifi_cell_type: Cell type label
timepoint: Time point of sample collection
fg: Foreground treatment condition
bg: Background treatment condition
pathway: Reactome Pathway ID
pathway_label: Reactome Pathway name
n_genes: number of genes in Hallmark pathway gene set
nomP: Nominal P-value reported by fastgsea
adjP: Adjusted P-value after multiple hypothesis testing correction
NES: Normalized Enrichment Score reported by fastgsea
leadingEdge: Semicolon-separated list of Leading Edge Genes reported by fastgsea

**Supplementary Table 8. Transcription Factor motif enrichment in differentially accessible peaks.** TF enrichment scores among up- and down-regulated DAPs computed using hypergeometric tests. Column descriptions:

aifi_cell_type: Cell type label
timepoint: Time point of sample collection
fg: Foreground treatment condition
bg: Background treatment condition
feature: Transcription factor motif ID from the CISBP database
nomP: Nominal P-value for hypergeometric test for enrichment reported by ArchR
adjP: Adjusted P-value for hypergeometric test for enrichment reported by ArchR
enrichment: Enrichment of foreground motif frequency relative to background motif frequency
(frac_fg_hits / frac_bg_hits)
n_fg_peaks: number of differentially accessible foreground peaks
n_fg_hits: number of foreground peaks containing a motif for the feature
frac_fg_hits: fraction of foreground peaks containing a motif for the feature
n_bg_peaks: number of matched background peaks
n_bg_hits: number of background peaks containing a motif for the feature
frac_bg_hits: fraction of background peaks containing a motif for the feature

**Supplementary Table 9. Differentially expressed gene sets from a literature review of Dexamethasone response in human cells.** DEG sets curated from previous studies of dexamethasone response in primary human cells and cell types. Column descriptions:

pathway: short pathway name for computational analyses
description: brief, human-readable description of gene set
display_label: short, readable label for use in display
cell_type: cell type used in experiment, if applicable
cell_line: specific cell culture line used in experiment, if applicable
direction: direction of gene expression change, up or down, caused by dexamethasone
experiment: Experimental method or platform
geo_accession: GEO Accession for raw data, if available
n_genes: Number of genes in gene set
genes: Semicolon-separated gene list
source_url: Web URL for publication or web site from which gene set is derived
source_year: Year of publication of the source for the gene set
source_section: Table or supplementary table number where gene set was found within source

**Supplementary Table 10. Intersections between differentially expressed gene sets.** Overlapping genes and hypergeometric test results for comparisons between DEG sets. Column descriptions:

aifi_cell_type: Cell type label
group1_treatment: Drug treatment in DEG set 1
group1_timepoint: Timepoint for DEG set 1
group1_direction: Direction of regulation for DEG set 1
group2_treatment: Drug treatment in DEG set 2
group2_timepoint: Timepoint for DEG set 2
group2_direction: Direction of regulation for DEG set 2
n_common: Number of common genes detected in the conditions for gene set 1 and gene set 2
n_ol: Number of overlapping genes between gene set 1 and gene set 2
nomP: Nominal P-value for hypergeometric test adjP: Adjusted P-value for hypergeometric test
l_genes: Semicolon-separated list of genes in the intersection between DEG set 1 and DEG set 2

**Supplementary Table 11. Mean Fluorescence Intensity changes in response to drug treatment combinations.** Changes to MFI of drug treatments compared to DMSO-only controls. Column descriptions:

subject: Subject ID
population: Gated flow cytometry population (CD4 or CD8)
treatment: Drug treatment or drug combination
timepoint: Time point of sample collection
feature: Flow cytometry feature
mfi: Mean Fluorescence Intensity (MFI) of feature
group_min: Minimum MFI value for the feature across all samples
shifted_mfi: MFI shifted based on minimum MFI to account for negative values
log_shifted_mfi: Log10 of the shifted MFI values
fold_change: Fold change (MFI_treat_ / MFI_DMSO_)
logFC: Log2 of fold change

**Supplementary Table 12. Activation marker expression in response to drug treatment combinations.** Counts, fractions, and CLR-transformed values for cells with or without activation marker expression (CD40L and CD69) in CD3/CD28 stimulation experiments. Column descriptions:

subject: Subject ID
treatment: Gated flow cytometry population (CD4 or CD8)
stimulation: Whether or not (yes/no) cells were stimulated with CD3/CD28
beads population: Gated flow cytometry population (CD4 or CD8)
feature: Flow cytometry feature
feature_gate: Pos/Mid/Neg gate based on feature expression
count: Number of cells in the feature gate
fraction: Fraction of cells in the feature gate
clr: Centered Log Ratio of fractions

**Supplementary Table 12.**
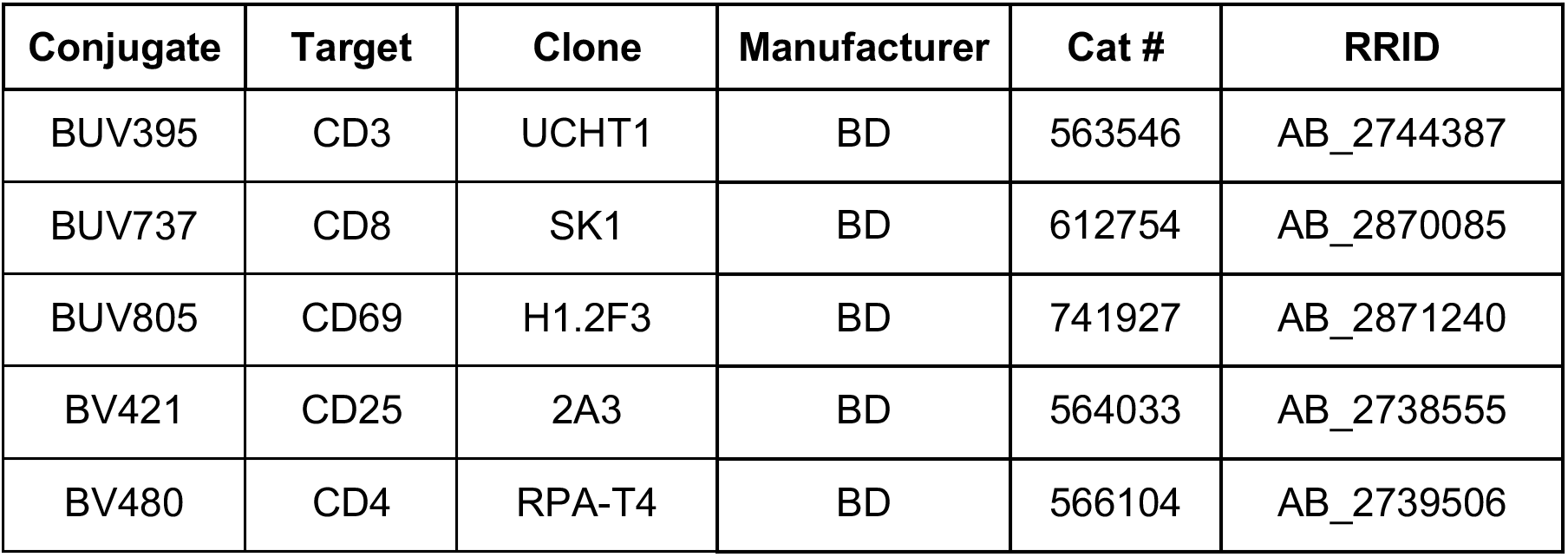
Surface Antibody Cocktail. List of antibodies used to quantify T cell populations and activation.

**Supplementary Table 13.** Intracellular Antibody Cocktail. List of antibodies used for intracellular flow cytometry to measure expression of Ikaros, Helios, and Aiolos.

| Conjugate | Target | Clone | Manufacturer | Cat # | RRID |
| --- | --- | --- | --- | --- | --- |
| eF450 | Granzyme B | M4TL33 | Invitrogen | 48-88960-42 | AB_2724392 |
| BV711 | Perforin | dG9 | BioLegend | 308130 | AB_2687190 |
| AF488 | Ikaros | R32-1149 | BD | 564867 | AB_2738990 |
| PE Dazzle 594 | Helios | 22F6 | BioLegend | 137232 | AB_2565797 |
| AF647 | Aiolos | EPR9342(B) | Abcam | ab198962 |  |

**Supplementary Table 14.** Single Color Controls. List of single-color control antibodies used for Flow Cytometry analysis.

| Conjugate | Target | Clone | Manufacturer | Cat # | RRID |
| --- | --- | --- | --- | --- | --- |
| eF450 | CD4 | SK3 | Invitrogen | 48-0047-41 | AB_1603230 |
| BV711 | CD3 | UCHT1 | BD | 563724 | AB_2744392 |
| AF488 | CD4 | SK3 | BioLegend | 344618 | AB_2228842 |
| PE Dazzle 594 | CD8 | SK1 | BioLegend | 344743 | AB_2566514 |
| AF647 | CD4 | SK3 | BioLegend | 344635 | AB_2566031 |

**Supplementary Table 15.**
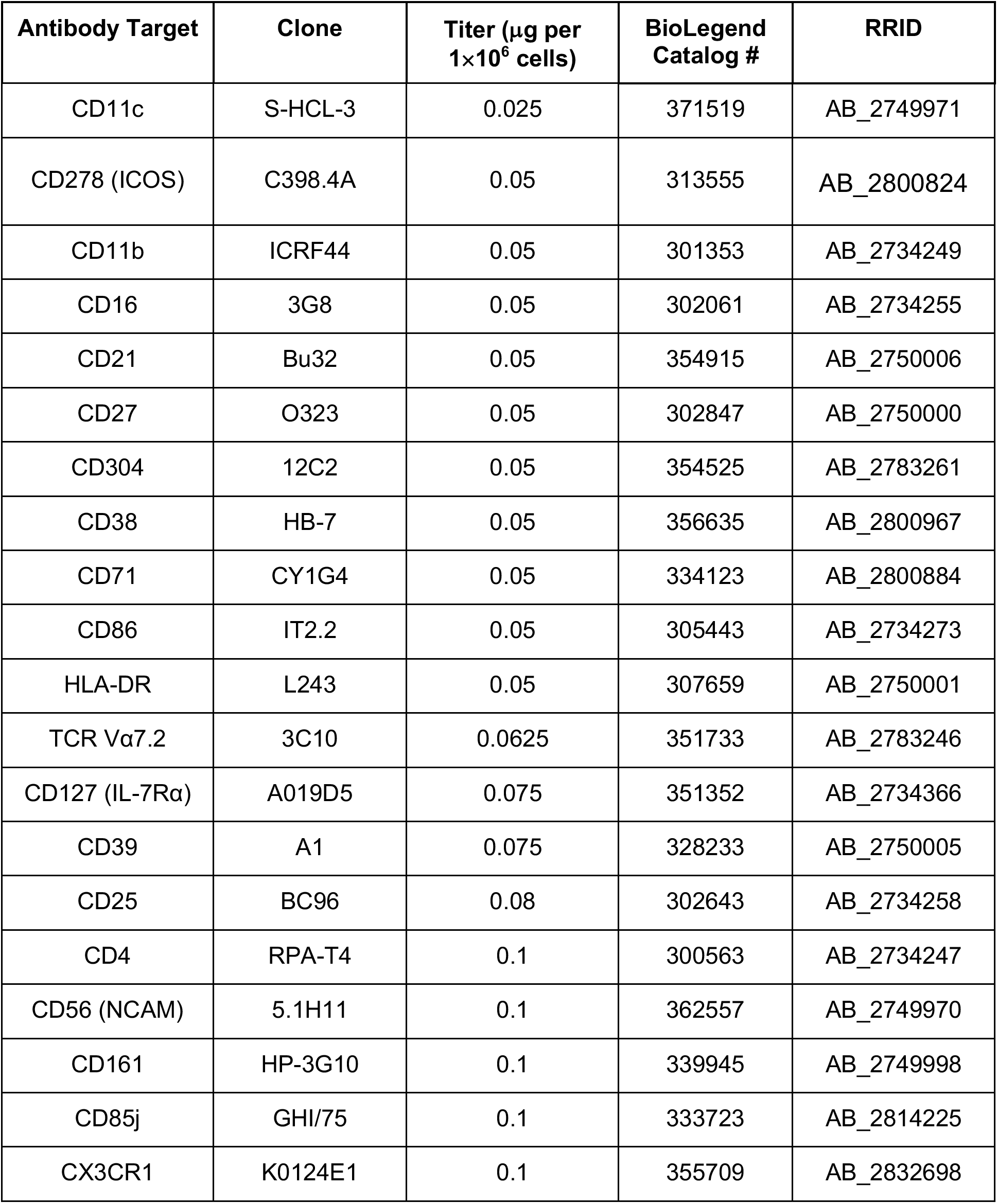
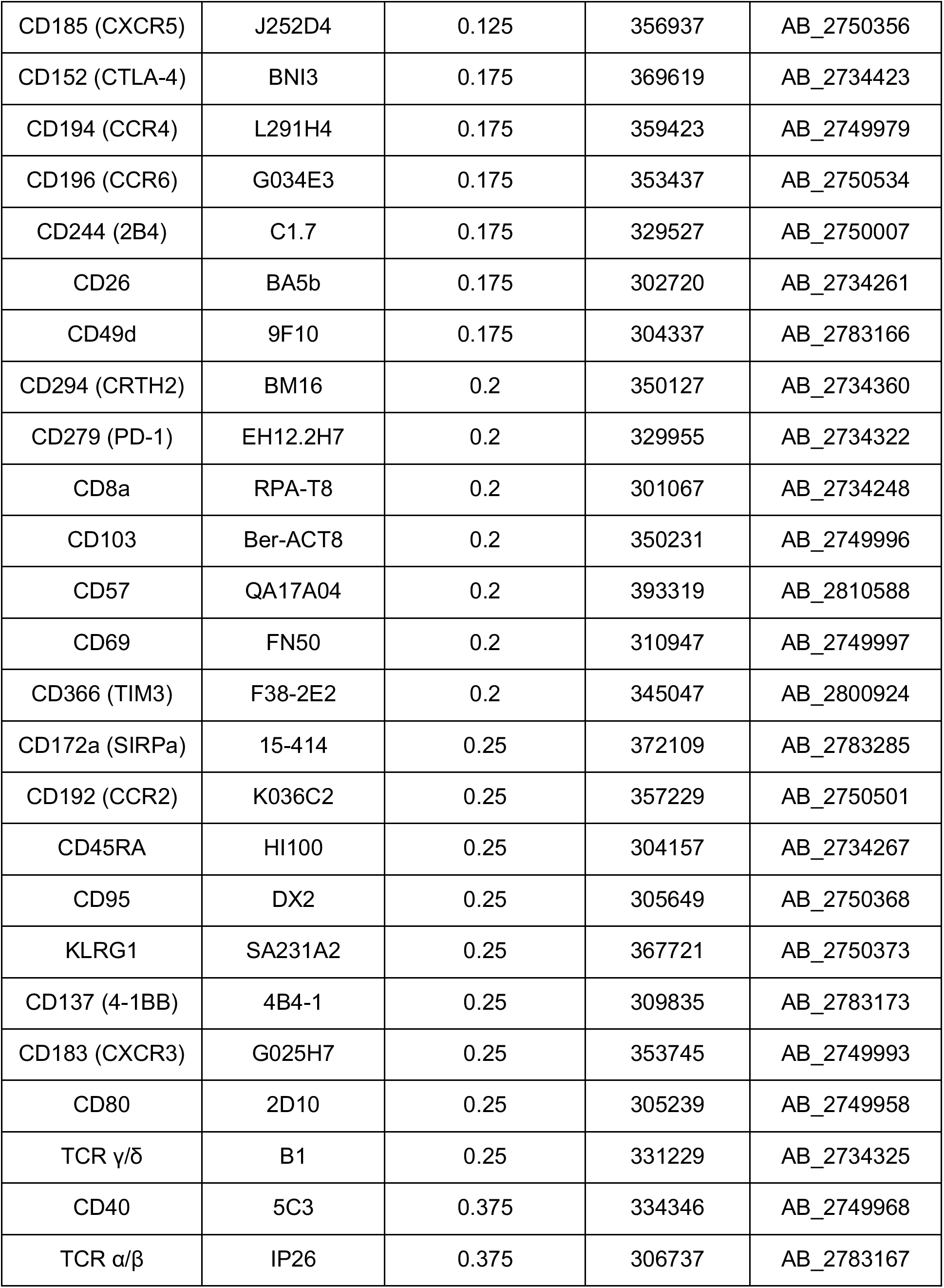
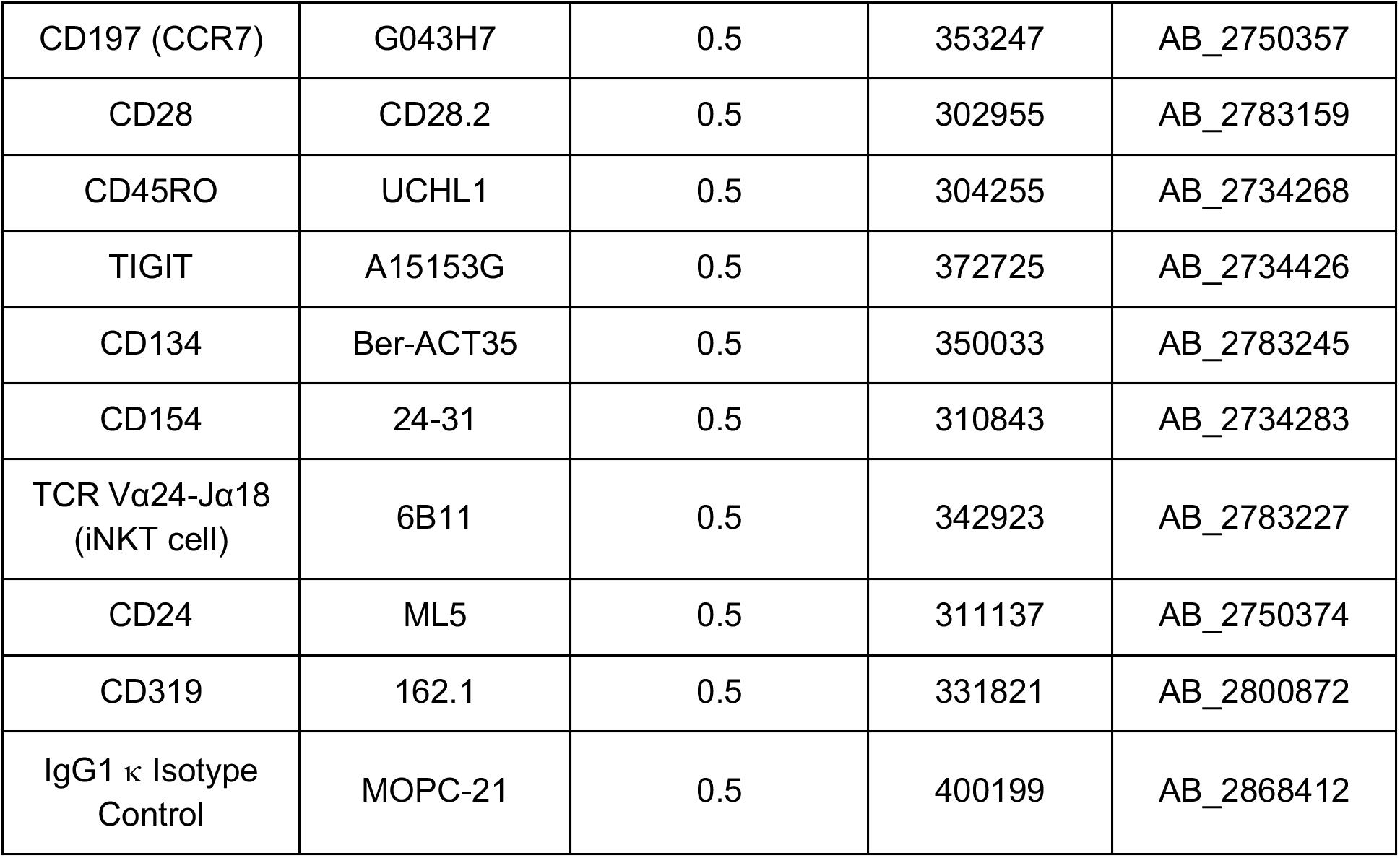
TEA-seq ADT Antibodies. List of antibodies used for antibody-derived tag (ADT) staining for TEA-seq. All antibodies listed are manufactered by BioLegend in the TotalSeq A format.

**Supplementary Table 16.** Cell Hashing Antibodies. List of antibodies used for cell hashing-based multiplexing staining for TEA-seq. All antibodies listed are manufactered by BioLegend in the TotalSeq A format.

| Antibody | Clone | Titer (μL per 1 M cells) | BioLegend Catalog # | RRID |
| --- | --- | --- | --- | --- |
| TotalSeq A0251 Hashtag 1 | LNH-94 2M2 | 2 | 394601 | AB_2750015 |
| TotalSeq A0252 Hashtag 2 | LNH-94 2M2 | 2 | 394603 | AB_2750016 |
| TotalSeq A0253 Hashtag 3 | LNH-94 2M2 | 2 | 394605 | AB_2750017 |
| TotalSeq A0254 Hashtag 4 | LNH-94 2M2 | 2 | 394607 | AB_2750018 |
| TotalSeq A0255 Hashtag 5 | LNH-94 2M2 | 2 | 394609 | AB_2750019 |
| TotalSeq A0256 Hashtag 6 | LNH-94 2M2 | 2 | 394611 | AB_2750020 |
| TotalSeq A0257 Hashtag 7 | LNH-94 2M2 | 2 | 394613 | AB_2750021 |
| TotalSeq A0258 Hashtag 8 | LNH-94 2M2 | 2 | 394615 | AB_2750022 |
| TotalSeq A0259 Hashtag 9 | LNH-94 2M2 | 2 | 394617 | AB_2750023 |
| TotalSeq A0260 Hashtag 10 | LNH-94 2M2 | 2 | 394619 | AB_2750024 |
| TotalSeq A0262 Hashtag 12 | LNH-94 2M2 | 2 | 394623 | AB_2750025 |
| TotalSeq A0263 Hashtag 13 | LNH-94 2M2 | 2 | 394625 | AB_2750026 |
| TotalSeq A0264 Hashtag 14 | LNH-94 2M2 | 2 | 394627 | AB_2750027 |

